# Structural basis of lipid head group entry to the Kennedy pathway by FLVCR1

**DOI:** 10.1101/2023.09.28.560019

**Authors:** Yeeun Son, Timothy C. Kenny, Artem Khan, Kıvanç Birsoy, Richard K. Hite

**Affiliations:** Structural Biology Program, Memorial Sloan Kettering Cancer Center, New York, NY, USA; BCMB Allied Program, Weill Cornell Graduate School, New York, NY, USA; Laboratory of Metabolic Regulation and Genetics, The Rockefeller University, New York, NY, USA

## Abstract

Phosphatidylcholine and phosphatidylethanolamine, the two most abundant phospholipids in mammalian cells, are synthesized *de novo* by the Kennedy pathway from choline and ethanolamine, respectively^1–6^. Despite the importance of these lipids, the mechanisms that enable the cellular uptake of choline and ethanolamine remain unknown. Here, we show that FLVCR1, whose mutation leads to the neurodegenerative syndrome PCARP^7–9^, transports extracellular choline and ethanolamine into cells for phosphorylation by downstream kinases to initiate the Kennedy pathway. Structures of FLVCR1 in the presence of choline and ethanolamine reveal that both metabolites bind to a common binding site comprised of aromatic and polar residues. Despite binding to a common site, the larger quaternary amine of choline interacts differently with FLVCR1 than does the primary amine of ethanolamine. Structure-guided mutagenesis identified residues that are critical for the transport of ethanolamine, while being dispensable for choline transport, enabling functional separation of the entry points into the two branches of the Kennedy pathway. Altogether, these studies reveal how FLCVR1 is a high-affinity metabolite transporter that serves as the common origin for phospholipid biosynthesis by two branches of the Kennedy pathway.

## Main

Phospholipids are integral components of membranes, whose amphipathic nature establishes barriers between cells and their environment and between subcellular compartments within cells. The two most common phospholipids in mammalian cells are phosphatidylcholine (PC) and phosphatidylethanolamine (PE), which together account for between 55-75% of the total phospholipid content^1,2^. PC and PE are synthesized *de novo* by the three-step Kennedy pathway from choline and ethanolamine via phospho– and cytidine-diphospho-intermediates^3–6^. Choline can also contribute to the biosynthesis of other lipids, one-carbon metabolism, and neurotransmitter synthesis^10^. We and others recently identified the ubiquitously expressed feline leukemia virus subgroup C cellular receptor 1 (FLVCR1) and its close paralog FLVCR2 as plasma membrane transporters that facilitate the uptake of extracellular choline into cells^11,12^. FLVCR1 is essential for viability in mice, and mice lacking functional FLVCR1 die around embryonic day E12.5^11^. In humans, several point mutations have been identified in *FLVCR1* that lead to posterior column ataxia and retinitis pigmentosa (PCARP), an autosomal-recessive neurodegenerative syndrome that is initiated during childhood and results in loss of vision, abnormal proprioception, and muscle atrophy^7–9^. Mutations in *FLVCR2* can lead to Fowler syndrome, a vascular disorder characterized by severe hydrocephaly, hypokinesia, and arthrogryposis^13–15^.

FLVCR1 is a member of the major facilitator superfamily (MFS) of membrane transporters that are characterized by possessing two bundles of six transmembrane helices (6TM) connected by an extended intracellular linker. Humans express over 100 MFS transporters that transport diverse substrates ranging from ions to sugars to porphyrins^16^. Structures of MFS transporters from different phyla resolved in inward-facing, occluded, and outward-facing states have revealed a conserved rocker-switch alternating access transport mechanism where the two 6TM bundles rock around a central substrate binding site to alternatively allow access from either the extracellular or intracellular side of the membrane^16–18^. Substrate transport by MFS transporters can be actively coupled to the symport or antiport of a second substrate, such as a proton, or can be passively coupled to electrochemical gradients^19^. For FLVCR1, the mechanisms of substrate recognition and transport are poorly understood. Here, we combine cryo-EM structural, biochemical, and metabolomic analyses to investigate the uptake of choline by FLCVR1. These studies unexpectedly revealed that FLVCR1 is also an ethanolamine transporter that can serve as the primary ethanolamine transporter in cells.

## FLVCR1 transports choline

To monitor choline transport by FLVCR1, we compared the uptake of radiolabeled choline ([methyl-^3^H]choline) by *FLVCR1*-knockout HEK293T cells or *FLVCR1*-knockout HEK293T cells expressing *FLCVR1* cDNA over the course of 30 minutes. Consistent with our previous results, we observe a time-dependent increase in choline uptake in cells expressing FLVCR1 that is severely impeded in the *FLVCR1*-knockout cells (**Fig. 1a**)^11^. To begin to understand the mechanisms of choline transport by FLVCR1, we first investigated the effects of monovalent and divalent cations on choline uptake by modifying the extracellular solution and measuring choline uptake after 30 minutes. Because cellular homeostasis is strongly influenced by the composition of the extracellular solution, we calculated the ratio of choline uptake by the *FLVCR1*-expressing cells at a particular condition to the uptake by the *FLVCR1*-knockout cells at the same condition. By comparing this ratio, we found that removal of sodium, potassium, calcium, or magnesium resulted in only minor changes in choline uptake by *FLVCR1*-expressing cells compared to *FLVCR1*-knockout cells, indicating that choline transport is not coupled to the influx of monovalent or divalent cations (**Extended Data** Fig. 1). We next examined the effect of extracellular pH, finding that acidic conditions inhibited choline uptake by FLVCR1. The inhibition followed a roughly inverse log-linear relationship from pH 4.2, where uptake by the *FLVCR1*-expressing cells was only slightly greater than uptake by the *FLVCR1*-knockout cells, to pH 9.2, where we observed maximal uptake compared to the *FLVCR1-*knockout cells. Together, these data indicate that choline transport by FLVCR1 is not accompanied by the symport of a proton, a monovalent cation or a divalent cation. We instead propose that FLVCR1 is a passive facilitator that takes advantage of the choline gradient between the extracellular solution, which has 5-10 µM choline in mammals^20–23^, and the cytosol, which has low choline concentrations due to the phosphorylation of choline by choline kinase alpha^24^.

**Fig. 1:**
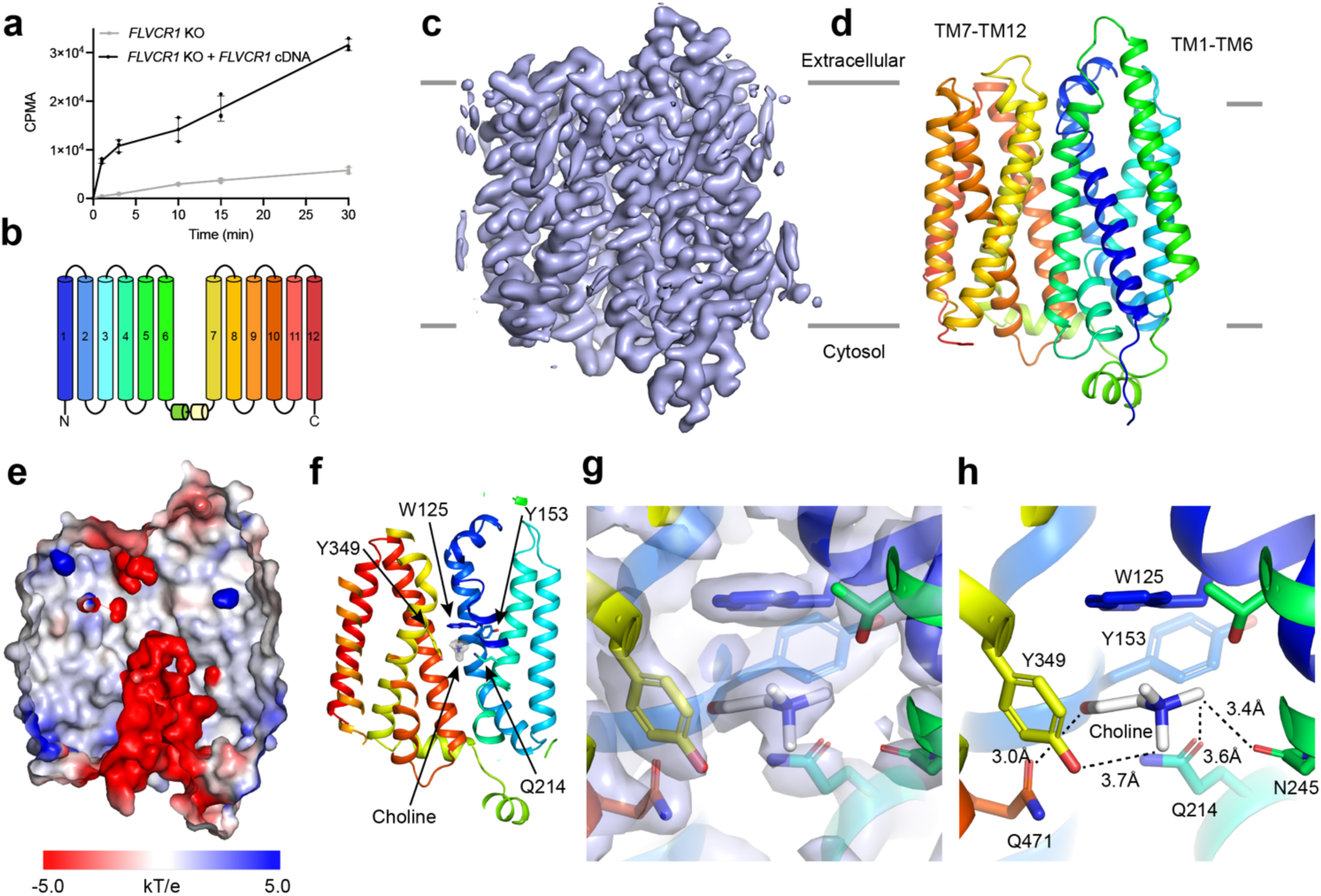
Structure of human FLVCR1 in a choline-bound state. **a**, Uptake of 21.5 μM choline by *FLVCR1*-knockout HEK293T cells expressing a vector control (grey) or *FLVCR1* cDNA (black) for 30 minutes. Error bars represent the standard deviation, n=3. **b**, Schematic of FLVCR1 colored by helix. **c-d**, Cryo-EM density map (c) and model of FLVCR1 (d). Model is colored by helix as in b. Grey lines correspond to the approximate position of the membrane. **e**, Central slice of FLVCR1 with surface colored by electrostatic potential. **f**, Central slice of FLVCR1 with choline and choline-coordinating residues shown as sticks. Density is shown as a grey isosurface and contoured at 3.0 σ. **g-h**, Choline-binding site. Residues that comprise the substrate-binding site are shown as sticks. Density is shown as a grey isosurface in (g) and contoured at 3.0 σ.

## Structure of human FLVCR1 in the presence of choline

To determine the mechanisms of choline recognition and transport, we collected cryo-EM images of detergent-solubilized human FLVCR1 in the presence of 1 mM choline chloride (**Extended Data** Fig. 2). Iterative classification of particles extracted from these images resulted in a single class whose reconstruction achieved a resolution of 2.60 Å that was suitable for building and refining a *de novo* model of human FLVCR1 with good geometry (**Fig. 1b-d**, **Extended Data Fig. 2 and Extended Data Table 1**). Similar to other members of the MFS superfamily, FLVCR1 is comprised of two 6TM domains that are connected by a flexible intracellular linker, with TM1-TM6 forming the N-terminal domain (NTD) and TM7-TM12 forming the C-terminal domain (CTD). The NTD and CTD are structurally homologous, with an RMSD of 3.5 Å, creating a pseudo-two-fold symmetry axis through the center of the transporter (**Extended Data** Fig. 3). A cavity extending from the cytosolic side of the transporter can be observed along this central axis between the two domains (**Fig. 1e**). This central cavity extends more than 30 Å into the transporter and is highly electronegative due to six aspartate and glutamate residues within and near the cavity (**Fig. 1e** **and Extended Data** Fig. 3). The minimum radius of the cavity is 2.1 Å, which may be wide enough to accommodate a choline molecule. The extracellular side of the cavity is sealed by TM1 and TM7, which adopt kinked conformations that bring them into direct contact near their extracellular ends (**Fig. 1f**).

Several non-protein densities can be observed in the cytosolic cavity, including one at the top of the central cavity that we assigned as a choline molecule based on its shape (**Fig. 1f**). The hydroxyl group of the modeled choline interacts with the side chains of Gln214 and Gln471, while the quaternary amine forms cation-ρε interactions with the side chains of Trp125 and Tyr349 (**Fig. 1g-h**). Also contributing to the binding site are Tyr153, Met154, and Asn245. The substrate-binding residues are conserved in FLVCR2, suggesting that the paralogs share a conserved substrate-binding site (**Extended Data** Fig. 4). This combination of aromatic and polar residues is reminiscent of the choline-binding sites in the prokaryotic periplasmic choline-binding protein ChoX from *S. meliloti* (PDB: 2REG) and the prokaryotic choline transporter LicB from *S. pneumoniae* (PDB: 7B0K), both of which are evolutionarily distinct from FLVCR1^25,26^.

We next performed a DALI search to identify homologous structures deposited in the Protein Data Bank, finding that the two most homologous structures are two prokaryotic MFS transporters in inward-facing states: *E. coli* SotB (PDB: 6KKL; RMSD = 2.3 Å), a drug-proton antiporter, and *E. coli* DgoT (PDB: 6E9N; RMSD = 2.6 Å), a D-galactonate transporter^27–29^ (**Extended Data** Fig. 3). Taken together with the open cytosolic cavity and the occupied substrate-binding site, we assign the structure determined in the presence of 1 mM choline as an inward-facing, choline-bound state.

## Structures of FLVCR1 in complex with endogenous ligands

For importers, an inward-facing, substrate-bound structure corresponds to a state following substrate binding and translocation from the extracellular space but prior to release into the cytosol. To gain additional insights into the transport mechanism, we next collected cryo-EM images of FLVCR1 in the absence of exogenous substrate. Iterative classification of the particle images resulted in two distinct states (**Fig. 2a,c, Extended Data** Figs. 2 and 5**, and Extended Data Table 1**). The first state, which was resolved at 2.63 Å, is nearly identical to the choline-bound structure (all-atom RMSD 0.3 Å) (**Extended Data** Fig. 5). Indeed, inspection of the substrate-binding site reveals that it is occupied by a non-protein density that matches the choline density that we resolved in the choline-bound state (**Fig. 2a-b**). Based on the overall similarity of these structures, as well as the similarity of their substrate-binding sites, we assigned the density as a bound choline. As no choline was added during the purification, the bound choline must have been purified along with the transporter, indicating that choline binds to the substrate-binding site of FLVCR1 with high affinity.

**Fig. 2:**
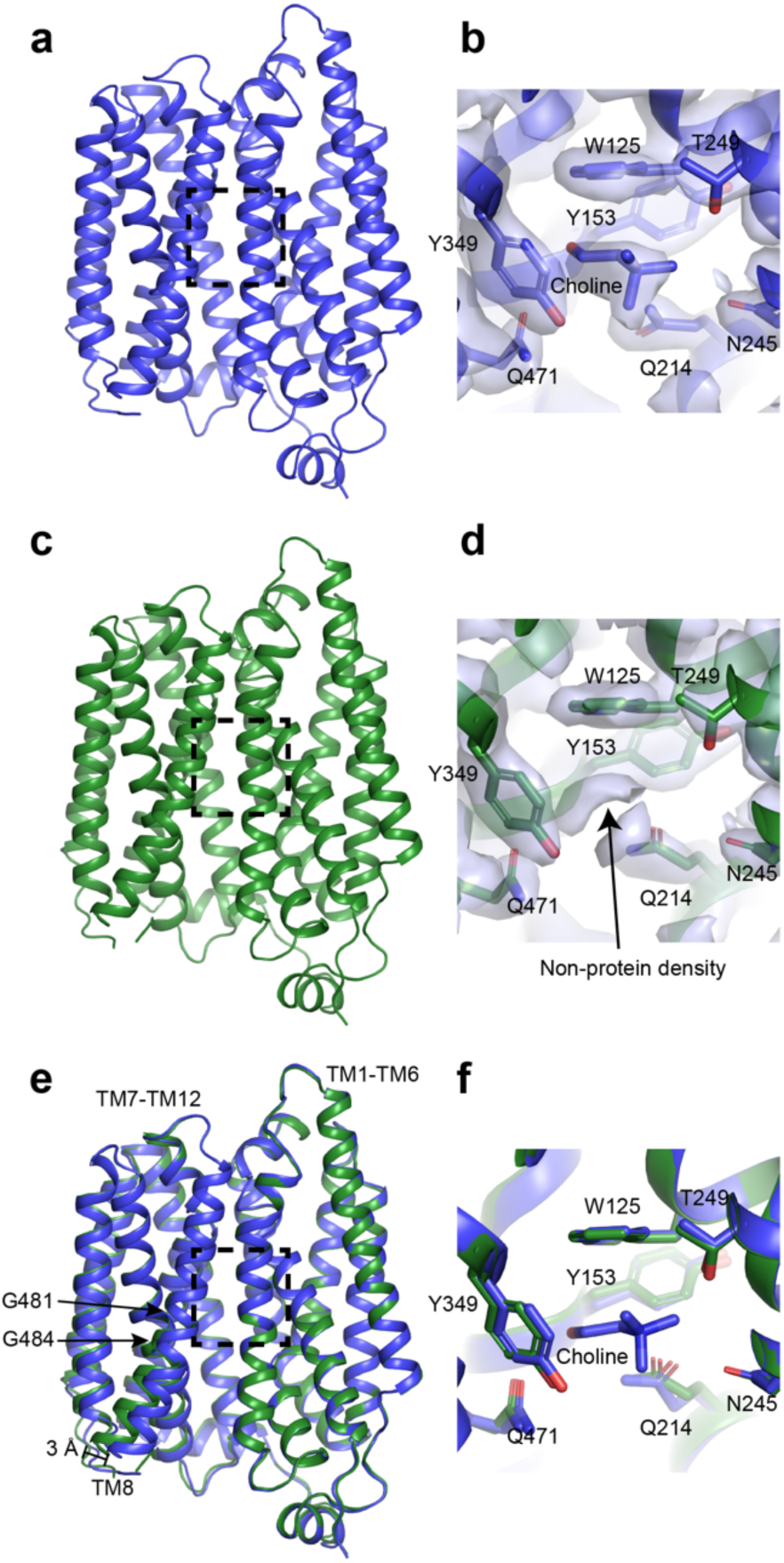
FLVCR1 bound to endogenous substrates. **a,c**, FLVCR1 in endogenous choline-bound (a) or endogenous ligand-bound (c) states, determined from particles imaged in the absence of exogenous substrate. Dashed box corresponds to the substrate-binding sites shown in b and d. **b,d**, Substrate-binding sites in endogenous choline-bound (b) or endogenous ligand-bound (d) states, determined from particles imaged in the absence of exogenous substrate. Residues and modelled substrates are shown as sticks. Density is shown as a grey isosurface and contoured at 3.0 σ. **e**, Superposition of endogenous choline-bound (blue) and endogenous ligand-bound (green) states. **f**, Superposition of substrate-binding sites in endogenous choline-bound (blue) and endogenous ligand-bound (green). Substrate-binding site residues and modelled substrates are shown as sticks.

The second state, which was resolved at 2.42 Å, also adopts an inward-facing state, but its conformation differs from the choline-bound states (**Fig. 2c,e**). The most pronounced difference is a tilt in TM8 that results in a 3 Å displacement of its cytosolic end, with Gly381 and Gly384 serving as the pivots (**Fig 2e**). Combined with more subtle movements of the cytosolic ends of the other helices in the CTD, the tilt in TM8 results in a widening of the central cavity in state 2 compared to the choline-bound states. Although the conformation of the substrate-binding site is similar to that of the choline-bound states, the density does not correspond to choline, nor to heme, which has been previously proposed to be a substrate of FLVCR1^30^ (**Fig. 2d,f**). The density is larger and extends deeper into the central cavity, indicating that one or more compounds other than choline were co-purified with transporter (**Fig. 2d**).

To generate hypotheses for additional substrates of FLVCR1, we used the Cancer Dependency Map (DepMap) project CRISPR 23Q2 public + score database ^31^ to identify genes that display co-essentiality with *FLVCR1*, as genes involved in the same metabolic pathways commonly exhibit similar patterns of essentiality^32,33^. Consistent with our previous report that *FLVCR1* serves a central role in choline metabolism^11^, the gene with the second strongest association with *FLVCR1* was *CHKA* (Pearson correlation = 0.48), which encodes choline kinase alpha that catalyzes the first step of the choline branch of the Kennedy pathway^31^ (**Fig. 3a-b**). Notably, the gene with the strongest association with *FLVCR1* was *ETNK1* (Pearson correlation = 0.51), which encodes ethanolamine kinase 1 that catalyzes the first step in the ethanolamine branch of the Kennedy pathway. Moreover, an association could also be identified between *FLVCR1* and *PCYT2* (Pearson correlation = 0.27), which encodes phosphate cytidylyltransferase 2 that catalyzes the second step of the ethanolamine branch of the Kennedy pathway. Together, these genetic associations suggest that in addition to its role in acquiring choline for the entry into the Kennedy pathway, FLVCR1 may also contribute to ethanolamine branch of the Kennedy pathway.

**Fig. 3:**
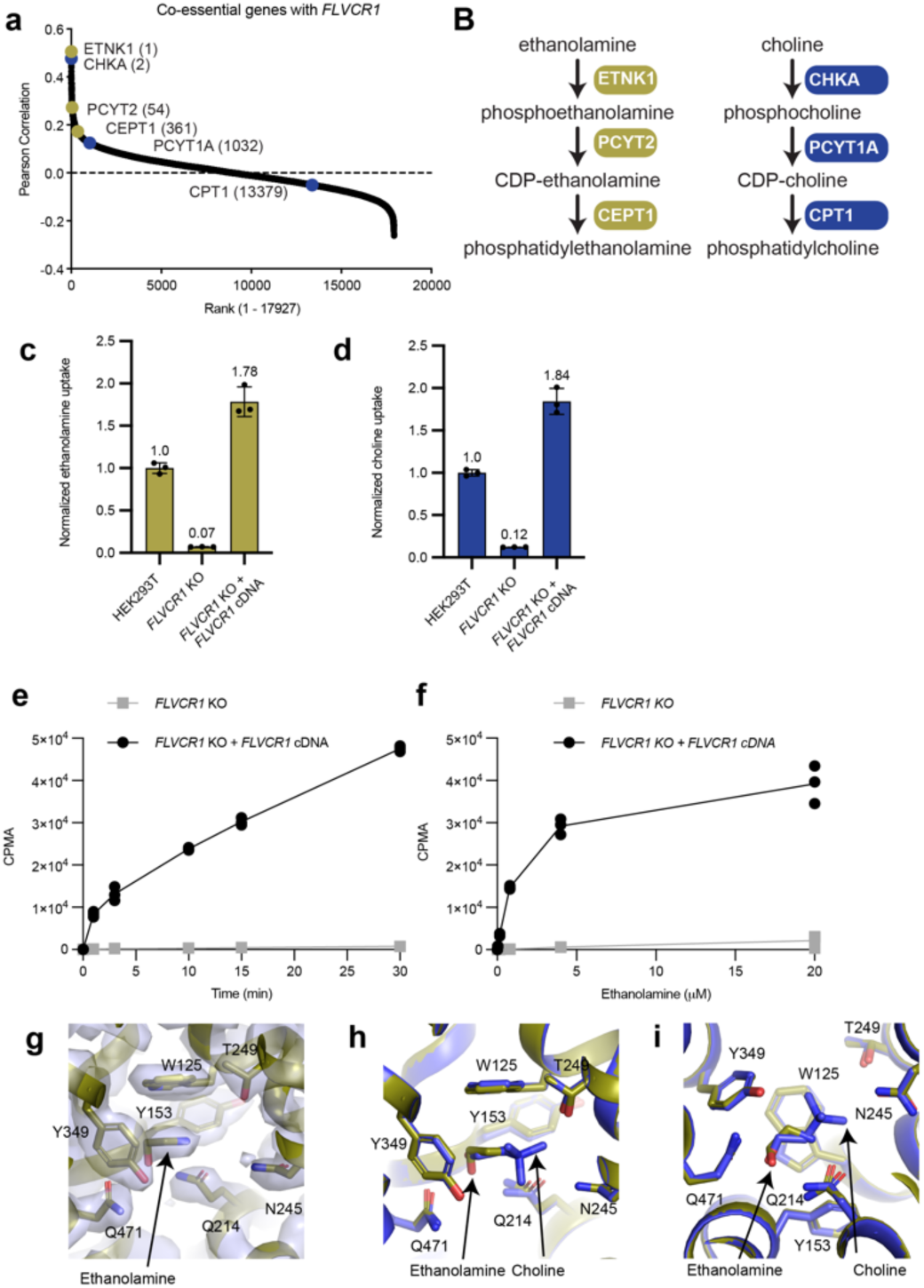
Ethanolamine and choline are transported by FLVCR1 through a common pathway. **a**, Pearson correlation of co-dependencies between *FLVCR1* and all genes computed from CRISPR DepMap Chronos 2023Q2. Genes in the ethanolamine (gold) and choline (blue) branches of the Kennedy pathway are highlighted. **b**, Schematic of the ethanolamine (left) and choline (right) branches of the Kennedy pathway. **c-d**, Uptake of 2 µM ethanolamine (c) or 21.5 µM choline (d) by control HEK293T cells, *FLVCR1*-knockout HEK293T cells expressing a vector control, and *FLVCR1*-knockout HEK293T cells expressing *FLVCR1* cDNA normalized to the uptake of control HEK293T cells. Error bars represent the standard deviation, n=3. **e**, Uptake of 2 µM ethanolamine by *FLVCR1*-knockout HEK293T cells expressing a vector control (grey) or *FLVCR1* cDNA (black), incubated for the indicated time point. Error bars represent the standard deviation, n=3. **f**, Uptake of ethanolamine by *FLVCR1*-knockout HEK293T cells expressing a vector control (grey) or *FLVCR1* cDNA (black), incubated with the indicated total ethanolamine concentration for 30 minutes. Error bars represent the standard deviation, n=3. **g**, Substrate-binding site in the ethanolamine-bound state. Residues and modelled substrates are shown as sticks. Density is shown as a grey isosurface and contoured at 3.0 α. **h-i**, Superposition of substrate-binding sites in the choline-bound (blue) and ethanolamine-bound (gold) states, shown in two views. Substrate-binding site residues and modelled substrates are shown as sticks.

## FLVCR1 is an ethanolamine transporter

We next tested whether FLVCR1 transports ethanolamine into cells by comparing the uptake of radiolabeled ethanolamine ([methyl-^3^H]ethanolamine) between HEK293T cells and *FLVCR1*-knockout HEK293T cells over the course of 30 minutes. Strikingly, the ability to take up ethanolamine dropped by 93% following FLVCR1 deletion, suggesting that FLVCR1 is the major route for ethanolamine uptake in this cell type (**Fig. 3c**). This decline in ethanolamine uptake was comparable—or even greater than— the impact of FLVCR1 deletion on choline uptake in the same cells (**Fig. 3d**). Furthermore, re-expression of *FLVCR1* cDNA in the *FLVCR1*-knockout cells enabled dose– and time-dependent uptake of ethanolamine into cells, confirming that FLVCR1 expression is limiting for ethanolamine and choline uptake in cultured cells (**Fig. 3e-f**).

To determine how FLVCR1 recognizes and transports ethanolamine, we next collected cryo-EM images of FLVCR1 in the presence of 1 mM ethanolamine. Following several rounds of classification, we resolved two inward-facing states at resolutions of 2.50 and 3.02 Å (**Extended Data** Figs. 2 and 6**, Extended Data Table 1**). The two states are highly similar, with TM8 adopting the straight conformation that we had observed in the choline-bound states (**Extended Data** Fig. 6). Indeed, the only remarkable differences between the two states determined in the presence of ethanolamine are the shapes of the densities occupying the substrate-binding site (**Fig. 3g-i** **and Extended Data** Fig. 6). In the lower-resolution reconstruction, the substrate-binding site density overlaps with the choline densities resolved in the choline-bound states, indicating that this state may represent another endogenous choline-bound state (**Extended Data** Fig. 6). Although the limited resolution precludes definitive assignment of the substrate, this structure suggests that high concentrations of ethanolamine may be insufficient to completely compete away endogenous choline from the substrate-binding site.

The substrate-binding site density of the higher-resolution reconstruction is smaller and well accommodates an ethanolamine molecule (**Fig. 3g**). We therefore assigned this as an ethanolamine-bound state. Notably, despite binding to a common site and not altering the global conformation of the transporter, ethanolamine interacts differently with FLVCR1 than does choline (**Fig. 3h-i** **and Extended Data** Fig. 6). Compared to choline, the entire ethanolamine is shifted by more than 1 Å within the substrate-binding pocket towards TM2. The shift positions ethanolamine deeper within in the aromatic pocket formed by Trp125, Tyr153 and Tyr349 whereas the quaternary amine of the larger choline is partially outside of the aromatic pocket. The shift deeper into the pocket enables the primary amine of ethanolamine to interact with the side chain of Gln214, which is too far away to directly interact with the quaternary amine of choline. Notably, Gln214 also interacts with the hydroxyl of ethanolamine in a manner similar to its interaction with the hydroxyl of choline. Thus, the primary amine of ethanolamine is coordinated by a combination of polar and cation-π interactions, whereas the quaternary amine of choline is exclusively coordinated by cation-π interactions. Moreover, the structures indicate that Gln214 serves a uniquely critical role in ethanolamine recognition by interacting with both its primary amine and its hydroxyl.

## Specificity of the substate-binding site

Although choline and ethanolamine can occupy a common substrate-binding site, they interact differently with FLVCR1 in the inward-facing state. To assess the role of these differential interactions on transport, we mutated Trp125, Tyr153, Gln214, Asn245, Tyr349 and Gln471 in the substrate-binding pocket to alanine, expressed the mutants in *FLVCR1*-knockout cells and measured uptake of radiolabeled ethanolamine or radiolabeled choline after 30 minutes. The W125A mutant reduced uptake of both ethanolamine and choline to the levels of the *FLVCR1*-knockout cells, indicating that Trp125 is essential for metabolite transport (**Fig. 4a-b**). The Y153A and Y349A mutants also greatly diminished the transport of ethanolamine and choline, indicating that the aromatic residues that comprise the substrate-binding site are all critical for transport. Notably, the reduction in ethanolamine uptake caused by the Y349A mutant was greater the effect on choline uptake, consistent with ethanolamine being more deeply embedded in the aromatic pocket of the substrate-binding site than choline. The Q214A mutant had an even more striking effect on transporter, with ethanolamine uptake being reduced by 94% compared to the wild-type transporter while choline uptake was unchanged. Mutation of the other two polar residues in the substrate-binding site, Asn245 and Gln471, had lesser effects on metabolite uptake, with the N245A mutant being slightly more severe. Thus, Gln214 serves a distinct role in ethanolamine uptake, likely due to its interaction with the charged primary amine. Collectively, these data demonstrate that although FLVCR1 transports ethanolamine and choline via a common pathway, the interactions that stabilize these metabolites differ due to their unique chemical properties.

**Fig. 4:**
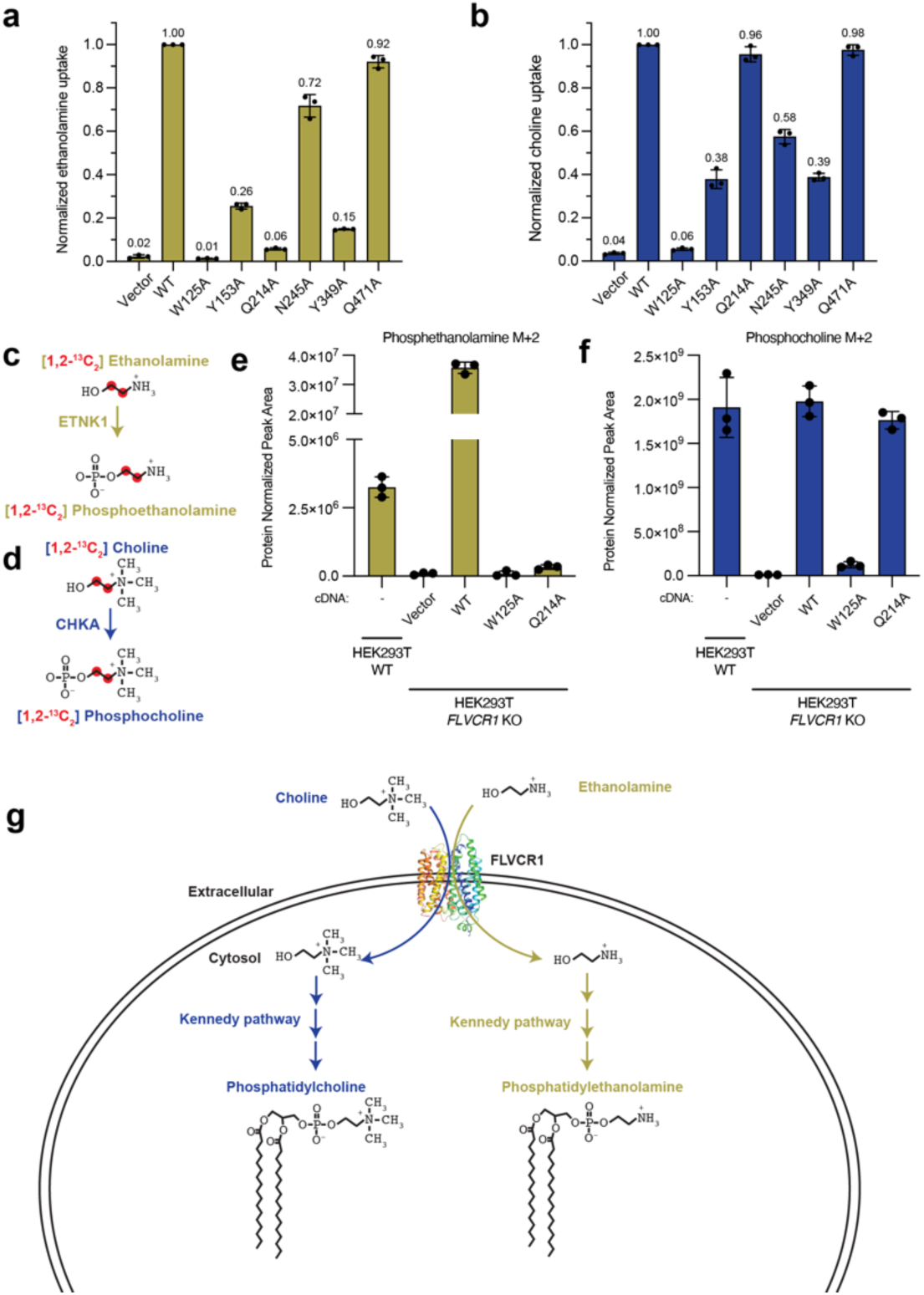
FLVCR1 fuels both branches of the Kennedy pathway. **a**, Uptake of 2 µM ethanolamine (a) or 2 µM choline (b) by *FLVCR1*-knockout HEK293T cells expressing a vector control or wild-type or mutant *FLVCR1* cDNA for 30 minutes normalized to the uptake by *FLVCR1*-knockout HEK293T cells expressing wild-type *FLVCR1*. Error bars represent the standard deviation, n=3. **c-d**, Schematic for tracing [1,2-^13^C_2_] ethanolamine (c) or [1,2-^13^C_2_] choline (d) into downstream metabolites. **e**, Abundance of phosphoethanolamine M+2 after incubation with 2 µM [1,2-^13^C_2_] ethanolamine for 1 hour in *FLVCR1*-knockout HEK293T cells expressing a vector control or wild-type or mutant *FLVCR1* cDNA. Data shown as mean ± standard deviation; n=3. **f**, Abundance of phosphocholine M+2 after incubation with 21.5 µM [1,2-^13^C_2_] choline for 1 hour in *FLVCR1*-knockout HEK293T cells expressing a vector control or wild-type or mutant *FLVCR1* cDNA. Data shown as mean ± standard deviation; n=3. **g**, Schematic of the phosphatidylcholine and phosphatidylethanolamine biosynthetic pathways.

We next examined whether ethanolamine and choline transport by FLVCR1 contributes to both branches of the Kennedy pathway in cells. We performed metabolite isotope tracing experiments using isotopically labeled choline ([1,2-^13^C_2_] choline) or isotopically labeled ethanolamine ([1,2-^13^C_2_] ethanolamine) in *FLVCR1*-knockout cells and those expressing wild-type and mutant *FLVCR1* (**Fig. 4c-f and Extended Data** Fig. 8). Consistent with our previous finding that *FLVCR1* is critical for the choline branch of the Kennedy pathway^11^, cells lacking *FLVCR1* displayed a 99% reduction in the incorporation of labeled choline into phosphocholine or the downstream metabolite betaine compared to the parental cells (**Fig. 4f** **and Extended Data** Fig. 8). Similarly, the loss of *FLVCR1* severely blunted the incorporation of labeled ethanolamine into phosphoethanolamine (**Fig. 4e**). The defects in incorporation of both choline and ethanolamine into downstream metabolites were specific to *FLVCR1* as they could be rescued by expression of the *FLVCR1* cDNA, but not the W125A loss-of-function mutant (**Fig. 4e-f**). Consistent with the specific requirement of Gln214 for ethanolamine transport by FLVCR1, incorporation of ethanolamine into phosphoethanolamine could not be rescued by the Q214A mutant, although the Q214A mutant was sufficient to rescue the incorporation of choline into phosphocholine and betaine. Thus, FLVCR1 is a choline and ethanolamine transporter that is responsible for their uptake to fuel both branches of the Kennedy pathway.

## Discussion

Phospholipid biosynthesis by the Kennedy pathway provides the phosphatidylcholine and phosphatidylethanolamine lipids that are essential for cellular proliferation. Here, we show that FLVCR1 is a transporter that serves as the common entry point for their biosynthesis, transporting extracellular choline and ethanolamine into the cytosol where they can be phosphorylated by choline kinase alpha and ethanolamine kinase 1, the enzymes that catalyze the first branches of the choline and ethanolamine branches of the Kennedy pathway, respectively (**Fig. 4g**). Several other transporters, including broadly-expressed CTL1 and CTL2, have been previously proposed to serve as ethanolamine transporters^34^. However, their reported affinities for ethanolamine are far below the concentrations present in plasma and we find that the loss of FLVCR1 largely eliminates ethanolamine uptake, suggesting that they are unlikely to serve as primary ethanolamine transporters.

Although FLVCR1 activity is regulated by extracellular pH, transport does not appear to be coupled to an exogenous energy source, suggesting that it is a passive facilitator. The activities of choline kinase alpha and ethanolamine kinase 1 ensure that cytosolic concentrations of ethanolamine and choline remain sufficiently low that there is a stable concentration gradient to drive their uptake from the extracellular space, a mechanism that is analogous to the uptake of sugars by the related GLUTS^35^.

FLVCR1 is a member of the ubiquitous MFS family of membrane transporters that utilize a conserved rocker-switch alternating access transport mechanism to move substrates across membranes^16^. When bound by choline or ethanolamine, FLVCR1 adopts a common inward-facing conformation that likely correspond to pre-release steps in the transport cycle. Notably, choline can co-purify with the transporter in these conformations, even when an ethanolamine is present at high concentrations for 30 minutes, suggesting that substrate release from the inward-facing state is very slow. An additional compound or compounds was also found to co-purify with FLVCR1 that stabilize a slightly different inward-facing state in which the central cavity is expanded, indicating that FLVCR1 can bind to multiple substrates and that these substrates can stabilize distinct states. However, as substrate release is a necessary step of the transport cycle, it is unclear if the slow off-rate from the resolved states is sufficient to enable choline and ethanolamine flux for entry into the Kennedy pathway and other biosynthetic reactions. Future studies will be needed to uncover the mechanisms of substrate release, as local or global conformational change may be needed to induce substrate release.

Despite binding to a common site, ethanolamine and choline interact differently with FLVCR1. Gln214 is notably required for ethanolamine transport, but not for choline transport. Tyr349 is also more important for uptake of ethanolamine than for choline. Future mutagenesis studies may uncover additional residues that can bias the substrate profile of the transporter. We recently demonstrated that the loss of FLVCR1 in mice leads to significant defects in mitochondrial metabolism and embryonic lethality around day E12.5, demonstrating that choline and/or ethanolamine transport by FLVCR1 for use in the Kennedy pathway is essential for mammalian viability^11^. Exploring the effects of mutations that alter the substrate profile of *FLVCR1* in cells and animals will enable a more complete understanding of the distinct physiological roles of ethanolamine and choline. Moreover, combining these substrate-specific mutations with variants in *FLVCR1* that lead to PCARP or variants in FLVCR2 that lead to Fowler syndrome, all of which are located outside of the substrate-binding site, may aid in understanding the molecular basis of disease (**Extended Data** Fig. 4)^7,8,13^.

## Methods

### Protein expression and purification

The gene encoding human FLVCR1 (hFLVCR1) was synthesized by Twist Biosciences and subcloned into a BacMam expression vector with a C-terminal mCerulean-tag fused via a short linker containing a PreScission protease site^36^. The plasmid was mixed 1:3 (w/w) with PEI 25 K (Polysciences, 23966-1) for 20 min at room temperature and then used to transfect HEK293S GnTi^-^ cells at 37°C in 5% CO2 (ATCC, CRL-3022). For a 1 L cell culture, 1 mg plasmid and 3 mg PEI 25 K were used. After 24 hr incubation at 37°C, valproic acid sodium salt (Sigma-Aldrich, P4543) was added to a final concentration of 2.2 mM, and cells were allowed to grow at 37°C for an additional 48 hr before harvesting. Cell pellets were washed in phosphate-buffered saline solution and flash frozen in liquid nitrogen. Expressed protein was solubilized in 2 % lauryl maltose neopentyl glycol (LMNG) (Anatrace, NG310), 0.2 % Cholesteryl Hemisuccinate Tris Salt (CHS) (Anatrace, CH210), 20 mM HEPES (pHed with KOH, pH 7.5), 150 mM KCl supplemented with protease-inhibitor cocktail (1 mM PMSF, 2.5 mg/mL aprotinin, 2.5 mg/mL leupeptin, 1 mg/mL pepstatin A) and DNase I. Solubilized proteins were separated by centrifugation 135,557 g for 50 min, followed by binding to anti-GFP nanobody resin for 3 hr. Anti-GFP nanobody affinity chromatography was performed by 3 column volumes of washing with SEC buffer containing 0.01 % LMNG, 0.001 % CHS, 20 mM HEPES (pHed with KOH, pH 7.5) and 150 mM KCl, followed by overnight PreScission digestion, and elution with wash buffer. Eluted protein sample was concentrated to a volume of 250 µl using CORNING SPIN-X concentrators (50 kDa cutoff) (Corning, 431490), followed by centrifugation 21,130 g for 15 min. Concentrated protein was further purified by size exclusion chromatography on a Superdex 200 Increase 10/300 GL (GE healthcare, GE28-9909-44) in SEC buffer. Peak fractions were pooled, mixed, and split into two samples in equal volume, of which one was incubated with choline chloride (Sigma-Aldrich, C7527) at a final concentration of 1 mM for 30 min on ice before concentrating to a protein concentration of 6 mg/ml, while the other was concentrated to a protein concentration of 6mg/ml without incubating with a ligand. Another hFLVCR1 expression and purification experiment was performed with identical procedures up to the size exclusion chromatography step, in which the SEC peak fractions were pooled, mixed, and incubated with ethanolamine (Sigma-Adrich, 398136) at a final concentration of 1 mM for 30 min on ice before concentrating to a protein concentration of 6 mg/ml.

### Electron microscopy sample preparation and data acquisition

3 μl of 6 mg/ml purified hFLVCR1 with incubated for 30 minutes with 1 mM choline chloride, or with ethanolamine chloride, or without exogenous substrate was applied to glow-discharged Au 400 mesh QUANTIFOIL R1.2/1.3 holey carbon grids (Quantifoil) and then plunged into liquid nitrogen-cooled liquid ethane with an FEI Vitrobot Mark IV (FEI Thermo Fisher). The sample was frozen at 4°C with 100 % humidity, using blotting times of 3.5 s to 4.5 s, blotting force of 0, and a waiting time of 10 s. Grids were either transferred to a 300 keV FEI Titan Krios microscopy equipped with a K3 summit direct electron detector (Gatan) or with a Falcon IV direct detector and a SeletrisX energy filter (ThermoFisher Scientific).

For images collected using a Gatan K3, the images were recorded with SerialEM^37^ in super-resolution mode at 29,000×, corresponding to super-resolution pixel size of 0.413 Å. Dose rate was 15 electrons/pixel/s, and the defocus range was −0.5 to −1.5 µm. Images were recorded for 3 s with 0.05 s subframes (total 60 subframes), corresponding to a total dose of 66 electrons/Å.

For images collected with a Falcon IV, the images were recorded with Leginon^38^ in pixel size of 0.725 Å. Dose rate was 5.67 electrons/pixel/s, and the defocus range was –0.5 to –1.5 μm. Images were recorded for 5 s with frame time of 0.003125 s, corresponding to a total dose of 53.98 electrons/ Å.

## Electron microscopy data processing

### Substrate-free sample, Gatan K3

5700 of sixty-frame super-resolution movies (0.413 Å/pixel) of hFLVCR1 without substrate incubation were collected using K3 summit direct electron detector (Gatan). The movies were gain corrected, Fourier cropped by two (0.826 Å/pixel) and aligned using whole-frame and local motion correction algorithms by cryoSPARC v3.2.0^39^. Blob-based autopicking in cryoSPARC v3.2.0 was implemented to select initial particle images, resulting in 5,926,467 particles^39^. Several rounds of 2D classification were performed and the best 2D classes were manually selected to generate the initial 3D model using the ab initio algorithm in cryoSPARC v3.2.0^39^. False-positive selections and contaminants were excluded through iterative rounds of heterogeneous classification using the model generated from the ab initio algorithm as well as several decoy classes generated from noise particles via ab initio reconstruction in cryoSPARC v3.2.0^39^, resulting in a stack of 2,709,623 particles. Then, 3D classification in cryoSPARC v3.2.0 ^40^ was performed using a focus mask that covers the FLVCR1 density but excludes the detergent micelle density, which we refer to as a FLVCR1 focus mask. The 3D classification did not resolve different conformations but separated the best-aligning particles from the rest of the particles that were seemingly in the same conformation, resulting in a stack of 824,310 best-aligning particles that yielded an improved reconstruction. After Bayesian polishing in Relion 3.1.2^41^, the polished particle stacks were refined using non-uniform refinement in cryoSPARC v3.2.0 with global CTF estimation, as well as higher order tetrafoil, anisotropic magnification, and aberration corrections, yielding a reconstruction of 2.80 Å. The particles were then classified via 3D classification in cryoSPARC v3.2.0 using the FLVCR1 focus mask^40^. The 3D classification did not resolve different conformations but separated the best-aligning particles from rest of the particles that were seemingly in the same conformation. The final reconstruction of 265,053 particles at resolution 2.66 Å was used for a reference for heterogenous classifications of other datasets described below.

### Choline sample, Falcon IV

3988 of one-thousand-six-hundred-frame movies of hFLVCR1 incubated with choline chloride were collected using a Falcon IV direct detector with a SelectrisX energy filter (ThermoFisher Scientific). The movies were up-sampled by 2, fractionated to 40, and aligned using whole-frame and local motion correction algorithms by cryoSPARC v4.2.1^39^. Blob-based autopicking in cryoSPARC was implemented to select initial particle images, resulting in 2,712,047 particles. False-positive selections and contaminants were excluded through iterative rounds of heterogeneous classification using the final reconstruction obtained from the substrate-free dataset collected with the Gatan K3 detector, as well as several decoy classes generated from noise particles via ab initio reconstruction in cryoSPARC v4.2.1^39^, resulting in a stack of 632,179 particles. After particle polishing in Relion 4.0.0^41^ with up-sampling factor of 2 and fractionation of 40, the particles were subjected to non-uniform refinement in cryoSPARC v4.2.1 with global CTF estimation as well as higher order tetrafoil, anisotropic magnification, and aberration corrections in cryoSPARC v4.2.1. The refined particles were then classified via 3D classification in cryoSPARC v4.2.1 using the FLVCR1 focus mask without additional angular alignment^40^. The 3D classification did not resolve different conformations but separated the best-aligning particles that adopted a single conformation. A non-uniform refinement followed by local-refinement of the subsequent 282,773 particles in cryoSPARC v4.2.1^39^ achieved a resolution of 2.60 Å. The final reconstruction was subjected to density modification using the two unfiltered half-maps with a soft mask in Phenix^42^, yielding an improved density map at 2.52 Å that we call choline-bound hFLVCR1. The density modified map was then used to build the choline-bound hFLVCR1 structure.

### Substrate-free sample, Falcon IV

9694 of one-thousand-six-hundred-frame movies of hFLVCR1 without substrate incubation were collected from two grids using a Falcon IV direct detector with a SelectrisX energy filter (ThermoFisher Scientific). The movies were up-sampled by 2, fractionated to 40, and aligned using whole-frame and local motion correction algorithms in cryoSPARC v4.2.1^39^. Blob-based autopicking in cryoSPARC was implemented to select initial particle images, resulting in stacks of 3,374,453 and 3,166,447 particles.

False-positive selections and contaminants were excluded through iterative rounds of heterogeneous classification using the final reconstruction obtained from the substrate-free dataset collected with K3 summit direct electron detector, as well as several decoy classes generated from noise particles via ab initio reconstruction in cryoSPARC v4.2.1^39^, resulting in stacks of 661,859 and 515,821 particles. After particle polishing in Relion 4.0.0.^41^, the stacks were subjected to non-uniform refinement in cryoSPARC v4.2.1 with global CTF estimation as well as higher order tetrafoil, anisotropic magnification, and aberration corrections in cryoSPARC v4.2.1. The two stacks were then classified via 3D classification in cryoSPARC v4.2.1^40^ using the FLVCR1 focus mask without additional angular alignment. Two classes of particles in distinct conformations were identified after the 3D classification. A non-uniform refinement followed by local refinement of one of the two classes resulted in reconstruction of 50,496 particles at 2.63 Å. Density modification of the first state using the two unfiltered half-maps with a soft mask in Phenix^42^ yielded an improved density map at 2.60 Å in which clear density for a choline was observed in the in the substrate-binding site and thus we call the endogenous choline-bound FLVCR1. A non-uniform refinement of the second class resulted in a reconstruction of 193,986 particles at 2.42 Å. Density modification of the second map using the two unfiltered half-maps with a soft mask in Phenix^42^ yielded an improved density map at 2.38 Å. Inspection of the second map revealed the presence of a density in the cavity that does not correspond to choline and we thus call this map the endogenous ligand-bound FLVCR1.

### Ethanolamine sample, Gatan K3

3675 of sixty-frame super-resolution movies (0.413 Å/pixel) of hFLVCR1 incubated with ethanolamine were collected using K3 summit direct electron detector (Gatan). The movies were gain corrected, Fourier cropped by two (0.826 Å/pixel) and aligned using whole-frame and local motion correction algorithms by cryoSPARC v3.2.0^39^. Blob-based autopicking in cryoSPARC v3.2.0 was implemented to select initial particle images, resulting in 5,046,088 particles^39^. False-positive selections and contaminants were excluded through iterative rounds of heterogeneous classification using the final reconstruction obtained from the substrate-free dataset collected with the Gatan K3 detector, as well as several decoy classes generated from noise particles via ab initio reconstruction in cryoSPARC v4.2.1^39^, resulting in a stack of 370,203 particles. After Bayesian polishing in Relion 3.1.2^41^, the polished particle stacks were refined using non-uniform refinement in cryoSPARC v3.2.0 with global CTF estimation, yielding a reconstruction of 2.65 Å. The refined particles were then classified via 3D classification in cryoSPARC v3.2.0 using the FLVCR1 focus mask^40^ without additional angular alignment^40^. Two classes of particles in distinct conformations were identified after the 3D classification. A non-uniform refinement of one of the two classes resulted in reconstruction of 36,781 particles at 3.02 Å that is equivalent to the choline-bound hFLVCR1 density map obtained from the movies of hFLVCR1 incubated with choline chloride collected with Falcon IV detector. A non-uniform refinement of the second class followed by local-refinement resulted in a reconstruction of 119,001 particles at 2.50 Å. Density modification of the first map using the two unfiltered half-maps with a soft mask in Phenix^42^ yielded an improved density map at 2.93 Å in which density corresponding to a bound choline could be observed in the substrate-binding site. Density modification of the second map using the two unfiltered half-maps with a soft mask in Phenix^42^ yielded an improved density map at 2.49 Å. Inspection of the map revealed the presence of a bound ethanolamine and we thus call this map the ethanolamine-bound FLVCR1.

### Model building and coordinate refinement

The structure of hFLVCR1 was automatically built into the choline-bound hFLVCR1 density map using ModelAngelo^43^. Densities corresponding to N– and C-terminus loops consisting of residues 1-98 and 517-555, respectively, were too poorly ordered and omitted from the model. The choline-bound hFLVCR1 model contains residues 99-516. The models were then manually rebuilt using COOT^44^ to fit the density. Choline and water molecules were modelled into non-protein density peaks using local geometry and a minimum threshold of 10.0 sigma in the density-modified and sharpened map. Atomic coordinates were refined against the density modified map using phenix.real_space_refinement with geometric and Ramachandran restraints maintained throughout ^45^.

The choline-bound model was docked into the endogenous choline map obtained from substrate-free sample using Chimera^46^. The model was then manually rebuilt using COOT ^44^ to fit the densityThe choline-bound model was docked into the endogenous choline map obtained from substrate-free sample using Chimera^45^. The model was then manually rebuilt using COOT ^46^ to fit the density. Densities corresponding to N– and C-terminus loops consisting of residues 1-98 and 518-555, respectively, were too poorly ordered and omitted from the model. The water molecules were modelled into non-protein density peaks using local geometry and a minimum threshold of 13.0 sigma in the density-modified and sharpened map. Atomic coordinates were refined against the density modified map using phenix.real_space_refinement with geometric and Ramachandran restraints maintained throughout.

Densities corresponding to N– and C-terminus loops consisting of residues 1-98 and 518-555, respectively, were too poorly ordered and omitted from the model. The water molecules were modelled into non-protein density peaks using local geometry and a minimum threshold of 13.0 sigma in the density-modified and sharpened map. Atomic coordinates were refined against the density modified map using phenix.real_space_refinement with geometric and Ramachandran restraints maintained throughout^45^.

The structure of unknown endogenous ligand-bound hFLVCR1 was initially built into the endogenous ligand-bound hFLVCR1 density map using ModelAngelo^43^. Densities corresponding to N– and C-terminus loops consisting of residues 1-96 and 519-555, respectively, were too poorly ordered and omitted from the model. The unknown endogenous ligand-bound hFLVCR1 model contains residues 97-518. The models were then manually rebuilt using COOT^46^The models were then manually rebuilt using COOT^44^ to fit the density. Ethanolamine and water molecules were modelled into non-protein density peaks using local geometry and a minimum threshold of 10.0 sigma in the density-modified and sharpened map. Atomic coordinates were refined against the density modified map using phenix.real_space_refinement with geometric and Ramachandran restraints maintained throughout ^45^.

The choline-bound model was docked into the ethanolamine-bound map using Chimera ^46^. Densities corresponding to N– and C-terminus loops consisting of residues 1-96 and 516-555, respectively, were too poorly ordered and omitted from the model. The ethanolamine-bound hFLVCR1 model contains residues 97-515. The model was then manually rebuilt using COOT ^44^ to fit the densityThe choline-bound model was docked into the ethanolamine-bound map using Chimera ^45^. Densities corresponding to N– and C-terminus loops consisting of residues 1-96 and 516-555, respectively, were too poorly ordered and omitted from the model. The ethanolamine-bound hFLVCR1 model contains residues 97-515. The model was then manually rebuilt using COOT ^46^ to fit the density. The ethanolamine and water molecules were modelled into non-protein density peaks using local geometry and a minimum threshold of 6.7 sigma in the density-modified and sharpened map. Atomic coordinates were refined against the density modified map using phenix.real_space_refinement with geometric and Ramachandran restraints maintained throughout. The ethanolamine and water molecules were modelled into non-protein density peaks using local geometry and a minimum threshold of 6.7 sigma in the density-modified and sharpened map. Atomic coordinates were refined against the density modified map using phenix.real_space_refinement with geometric and Ramachandran restraints maintained throughout ^45^.

The choline-bound model was docked into the endogenous choline map obtained from ethanolamine-incubated sample using Chimera^46^. Densities corresponding to N– and C-terminus loops consisting of residues 1-98 and 514-555, respectively, were too poorly ordered and omitted from the model. The ethanolamine-bound hFLVCR1 model contains residues 99-513. The model was then manually rebuilt using COOT^44^ to fit the density. The ethanolamine and water molecules were modelled into non-protein density peaks using local geometry and a minimum threshold of 8.0 sigma in the density-modified and sharpened map. Atomic coordinates were refined against the density modified map using phenix.real_space_refinement with geometric and Ramachandran restraints maintained throughout ^45^.

### Fluorescence size exclusion chromatography (FSEC)

Mutant hFLVCR1 constructs were generated via PCR using the WT hFLVCR1 construct, the primers listed in ‘Oligonucleotide sequences’ (synthesized by IDT) and Q5 polymerase (New England Biolabs, Q5 high fidelity 2X master mix M0492S), followed by HiFi-DNA assembly (New England Biolabs, E2621L) using the BacMam vector^36^ linearized with XhoI and EcoRI. All construct sequences were validated by Sanger sequencing.

*FLVCR1*-knockout HEK293T cells^11^ were seeded at a concentration of 500,000 cells per 6-well plate. The following day, the plasmid encoding mCerulean-tagged WT or mutant hFLVCR1 was mixed 1:3 (w/w) with PEI 25 k (Polysciences, Inc) for 20 min at room temperature and then used to transfect the *FLVCR1*-knockout HEK293T cells. After 24 hr incubation at 37 °C, valproic acid sodium salt (Sigma-Aldrich, P4543) was added to a final concentration of 2.2 mM, and cells were allowed to grow at 37 C°for an additional 48 hr before harvesting. Cell pellets were washed in phosphate-buffered saline solution and flash frozen in liquid nitrogen. Expressed proteins were solubilized in 2 % n-Dodecyl-β-D-maltoside (DDM) (Anatrace, D310), 20 mM HEPES pH 7.5 (pHed with KOH, pH 7.5), 150 mM KCl supplemented with protease-inhibitor cocktail (1 mM PMSF, 2.5 mg/mL aprotinin, 2.5 mg/mL leupeptin, 1 mg/mL pepstatin A) and DNase I (Worthington Biochemical, LS002139). Solubilized proteins were separated by centrifugation 15,871 g for 45 min. Separated proteins were injected to and monitored by fluorescence size exclusion chromatography on a Superose 6 Increase 10/300 GL (GE healthcare, GE29-0915-96) in a buffer composed of 1 mM DDM, 150 mM KCl, 20 mM HEPES pH 7.5 (pHed with KOH, pH 7.5). Fluorescence was monitored at 433/475 nm for excitation/emission wavelength, respectively.

### Generation of stable cell lines

Genes encoding WT or mutant hFLVCR1 in the BacMam vector^36^ were subcloned into pMXS-IRES-BLAST (Cell Biolabs, RTV-016) linearized with BamHI and NotI using Gibson Assembly (New England Biolabs, E2611) ^11^ using another set of primers listed in “Oligonucleotide sequences.” All construct sequences were validated by Sanger Sequencing. The cDNA expressing vectors along with retroviral packaging vectors Gag-Pol and VSG-G were transfected into HEK293T cells using X-tremeGENE 9 DNA Transfection reagent (Roche, 6364787001). Virus-containing supernatant was collected 48 hr after transfection and passed through a 0.45 μm filter. *FLVCR1*-knockout HEK293T cells^11^ in 6-well tissue culture plates were spin-infected with virus and 4 μg/mL polybrene by centrifugation at 2,200 rpm for 80 min. Cells were selected by 20 μg/mL blasticidin (Invivogen, ant-bl-1). For all constructs the matching vector without insert was used as a control. All cell lines were grown in RPMI 1640 medium (Gibco, 11875-093) containing 2 mM glutamine supplemented with 10 % fetal bovine serum (FBS) and were maintained at 37°C and 5 % CO2.

### Radioactive substrate uptake experiments

Radioactive substrate uptake experiments were performed as in^11^ unless otherwise stated. Briefly, cells were seeded in RPMI 1640 medium (Gibco, 11875-093) containing 2mM glutamine supplemented with 10 % fetal bovine serum (FBS) at 250,000 per mL or 500,000 cells per well in a 6-well plate in triplicate. The following day, media was aspirated and cells were incubated in room temperature Krebs-Ringer Buffer (Alfa Aesar, J67795) for 30 minutes. Cells were then incubated with indicated concentration of choline chloride or ethanolamine in room temperature Krebs-Ringer Buffer for timepoints described in figure legends. For all concentrations of choline chloride and ethanolamine, 0.093% of the total choline or ethanolamine concentration stated on figure legends was radioactive ([Methyl-^3^H]-Choline Chloride; Perkin Elmer, NET109001MC) (Choline chloride; Sigma-Aldrich, C7527) (ethanolamine [1-^3^H] hydrochloride; American Radiolabeled Chemicals, ARC-0216A) (ethanolamine; Sigma-Aldrich, 398136). For example, 20 nM was radioactive of total 21.5 μM choline used, and 1.86 nM was radioactive of total 2 μM ethanolamine used. Following incubation, cells were washed twice with ice cold Krebs-Ringer Buffer on ice. Cells were then solubilized with 200 μL of 1 % SDS 0.2 N NaOH and transferred to a scintillation vial with 10 mL of Insta-Gel Plus scintillation cocktail (Perkin Elmer, 601339). Radioactivity was measured with a liquid scintillation analyzer (Perkin Elmer, Tri-Carb 2910-TR).

For some experiments, Krebs-Ringer Buffer without sodium (125 mM KCl 2 mM CaCl_2_, 1 mM MgCl_2_, 25 mM KHCO_3_, 5.5 mM HEPES, 1 mM D-glucose; pH 7.2 with KOH/HCl), or without potassium (125 mM NaCl, 2 mM CaCl_2_, 1 mM MgCl_2_, 25 mM NaHCO_3_, 5.5 mM HEPES, 1 mM D-glucose; pH 7.2 with NaOH/HCl), or without calcium (120 mM NaCl, 5 mM KCl, 3 mM MgCl_2_, 25 mM NaHCO_3_, 5.5 mM HEPES, 1 mM D-glucose; pH 7.2 with NaOH/HCl), or without magnesium (120 mM NaCl, 5 mM KCl, 3 mM CaCl_2_, 25 mM NaHCO_3_, 5.5 mM HEPES, 1 mM D-glucose; pH 7.2 with NaOH/HCl) were used. For some experiments, Krebs-Ringer buffer with its pH adjusted to 4.2 5.2, 6.2, 7.2, 8.2, 9.2, and 10.2 using NaOH or HCl were used. In these experiments, the same Krebs-Ringer Buffer was used for incubation and washings.

## FLVCR1 Coessentiality Analysis

To perform a coessentiality mapping, we analyzed the DepMap 23Q2 Public+Score, Chronos dataset that contains the genetic perturbation scores of 1,095 cancer cell lines^31^. Specifically, we computed Pearson correlation values in a pairwise manner between the perturbation scores of FLVCR1 and 17,927 other genes present in the dataset. The code used to perform the analysis is available on Github (https://github.com/artemkhan/Coessentiality_DepMAP_FLVCR1.git).

## Immunoblotting

Cells were lysed in RIPA buffer (Cell Signaling Technology, 9806S) supplemented with protease-inhibitor cocktail (1 mM PMSF, 2.5 mg/mL aprotinin, 2.5 mg/mL leupeptin, 1 mg/mL pepstatin A) and phosphatase inhibitor cocktail (Sigma-Aldrich, P0044). Lysates were centrifuged at 18,000 g for 10 min. Total protein was quantified using BCA <u>Protein Assay</u> Kit (Thermo Fisher, 23227) with provided albumin standard used as a protein standard. Samples were resolved on 4-12% Bis-Tris gels (Thermo Scientific, NW04125BOX) and analyzed by standard immunoblotting techniques. Briefly, gels were transferred in Transfer buffer (25 mM Tris base, 190 mM glycine, 20 % methanol, pH 8.3) to nitrocellulose membranes (Bio-Rad, 1620115) and incubated with primary antibodies (FLVCR, Santa Cruz Biotechnology sc-390100; GAPDH, Cell Signalling Technology 5174) at 4°C overnight.

Secondary antibody incubation was performed at room temperature for 1 hr using anti-mouse IgG-HRP (Santa Cruz Bioctechnology, sc-525408) and anti-rabbit IgG-HRP (Cell Signalling Technology #7074). Washes were performed with 0.1% Tween-20 tris buffered saline and blots developed using ECL western blotting detection reagents (Cytiva, RPN2209).

## Isotope tracing

Polar metabolomics and isotope tracing experiments were performed as previously described^11^. Briefly, 250,000 cells were seeded per 6-well in triplicate per condition. For choline tracing experiments, cells were seeded in choline depleted medium and the next day changed to fresh choline depleted medium supplemented with 21.5 uM [1,2,-^13^C^2^]Choline chloride (Cambridge Isotope Laboratories, CLM-548.01). For ethanolamine tracing experiments, cells were seeded in RPMI 1640 medium (Gibco, 11875-093) with 2 mM glutamine supplemented with 10 % dialyzed fetal bovine serum (Gibco, 26400-044) and 1 % penicillin streptomycin and the next day changed to fresh medium with 2 uM [1,2,-^13^C^2^]ethanolamine hydrochloride (Cambridge Isotope Laboratories, CLM-274-0.1). For both choline and ethanolamine tracing, incubation was performed for 1 hour in a tissue culture incubator. Polar metabolites were extracted and analyzed by LC/MS as previously reported^11^.

## Figures

Figures were prepared with PyMol (www.pymol.org), ChimeraX ^47^, CAVER^48^, GraphPad Prism (www.graphpad.com), Clustal Omega^49^, and pyBoxshade (https://github.com/mdbaron42/pyBoxshade).

## Oligonucleotide sequences

**Primers used to clone from BacMam vector to pMXS-IRES-BLAST vector**

5’ GGGCGGAATTTACGTAGCCTAGATAGCACTCTCAGATTGTTTACTC 3’

5’ GCCGGATCTAGCTAGTTAATTAAATGGCCCGCCCCG 3’

## WT hFLVCR1

Forward primer used for cloning synthesized WT hFLVCR1 into BacMam expression vector

GCTAGCCTCGAGCCACCATGGCCCGCCCCGATGACGA

Reverse primer used for cloning synthesized WT hFLVCR1 into BacMam expression vector

TCCAAAGAATTCGAGATAGCACTCTCAGATTGTT

cDNA (Codon Optimized; 5’-3’)

ATGGCCCGCCCCGATGACGAAGAGGGAGCTGCCGTAGCTCCTGGCCATCCACTGGCCAAGGGGTATCTGCCTCTCCCACGGGGTGCCCCTGTGGGAAAAGAATCCGTCGAACTTCAAAATGGCCCTAAGGCCGGAACTTTTCCTGTTAACGGCGCGCCTCGTGATTCTCTGGCAGCTGCTAGCGGTGTATTGGGTGGACCACAAACCCCACTTGCACCCGAAGAAGAAACACAAGCCAGATTGCTGCCAGCCGGCGCGGGCGCCGAAACCCCCGGCGCCGAATCATCACCACTTCCTCTGACAGCCCTGTCTCCTAGACGATTTGTCGTCCTGCTCATATTTAGTCTTTACTCTCTCGTGAACGCATTCCAATGGATACAGTATTCAATAATCTCAAATGTGTTTGAAGGATTTTATGGCGTGACTCTGTTGCATATAGATTGGCTTAGCATGGTCTATATGCTCGCTTATGTTCCTCTGATATTTCCAGCTACATGGCTTCTCGATACTAGGGGTCTCCGCCTTACAGCACTCCTCGGGTCAGGTCTGAATTGTCTCGGAGCATGGATAAAATGTGGATCCGTCCAACAACACCTGTTTTGGGTGACGATGCTCGGGCAATGTCTTTGTTCAGTTGCACAAGTCTTTATTCTCGGGCTGCCATCAAGAATTGCTTCTGTCTGGTTCGGCCCTAAGGAAGTAAGCACGGCCTGCGCAACGGCAGTCCTTGGGAACCAACTTGGCACCGCTGTCGGGTTCCTGCTGCCACCCGTGCTGGTGCCTAATACCCAAAACGATACGAACCTGCTCGCCTGCAACATTTCCACTATGTTCTACGGCACCTCTGCAGTGGCGACTCTCCTGTTCATACTGACCGCGATCGCGTTCAAGGAGAAGCCGCGCTACCCACCGTCACAAGCCCAGGCTGCCTTGCAGGATTCACCACCCGAGGAATATAGCTACAAGAAGAGCATCAGGAATTTGTTCAAGAATATCCCGTTCGTGCTGTTGCTGATTACCTACGGCATTATGACAGGGGCATTTTATAGTGTGTCAACTCTCCTCAATCAAATGATTCTCACATATTATGAAGGTGAGGAAGTGAACGCAGGCAGAATCGGCCTGACCCTTGTGGTCGCCGGCATGGTAGGTAGCATCCTGTGCGGTTTGTGGCTCGACTACACCAAGACGTATAAGCAAACGACCCTTATTGTCTACATACTGTCCTTCATCGGCATGGTCATTTTCACCTTTACTCTGGATCTGAGGTACATCATTATTGTCTTCGTGACCGGTGGCGTCCTGGGGTTCTTTATGACAGGCTATCTGCCCCTGGGGTTCGAGTTCGCCGTCGAGATTACATATCCCGAGTCAGAGGGGACATCTTCCGGGCTGCTGAACGCAAGCGCCCAAATTTTCGGTATCCTGTTTACCCTGGCCCAGGGGAAATTGACCTCTGATTACGGACCGAAAGCTGGTAATATCTTCCTTTGCGTGTGGATGTTCATTGGGATTATCCTGACGGCCCTGATTAAATCCGACCTTCGGCGGCATAATATCAACATTGGGATCACCAACGTCGACGTCAAGGCCATCCCCGCCGATAGCCCGACTGATCAGGAGCCTAAGACTGTGATGCTGAGTAAACAATCTGAGAGTGCTATCTAG

## W125A hFLVCR1

**Forward primer 1** (5’-3’)

GTCCGAAGCGCGCTAGC

**Reverse primer 1** (5’-3’)

GAATACTGTATGGCTTGGAATGCGTTC

**Forward primer 2** (5’-3’)

GAACGCATTCCAAGCCATACAGTATTC

**Reverse primer 2** (5’-3’)

GGACCTTGAAACAAAACTTCCAAAGAATTCGAG

**cDNA** (Codon Optimized; 5’-3’)

ATGGCCCGCCCCGATGACGAAGAGGGAGCTGCCGTAGCTCCTGGCCATCCACTGGCCAAGGGGTATCTGCCTCTCCCACGGGGTGCCCCTGTGGGAAAAGAATCCGTCGAACTTCAAAATGGCCCTAAGGCCGGAACTTTTCCTGTTAACGGCGCGCCTCGTGATTCTCTGGCAGCTGCTAGCGGTGTATTGGGTGGACCACAAACCCCACTTGCACCCGAAGAAGAAACACAAGCCAGATTGCTGCCAGCCGGCGCGGGCGCCGAAACCCCCGGCGCCGAATCATCACCACTTCCTCTGACAGCCCTGTCTCCTAGACGATTTGTCGTCCTGCTCATATTTAGTCTTTACTCTCTCGTGAACGCAT TCCAA**GCC**ATACAGTATTCAATAATCTCAAATGTGTTTGAAGGATTTTATGGCGTGACTCTG TTGCATATAGATTGGCTTAGCATGGTCTATATGCTCGCTTATGTTCCTCTGATATTTCCAGCT ACATGGCTTCTCGATACTAGGGGTCTCCGCCTTACAGCACTCCTCGGGTCAGGTCTGAATTGTCTCGGAGCATGGATAAAATGTGGATCCGTCCAACAACACCTGTTTTGGGTGACGATGCTCGGGCAATGTCTTTGTTCAGTTGCACAAGTCTTTATTCTCGGGCTGCCATCAAGAATTGCTTCTGTCTGGTTCGGCCCTAAGGAAGTAAGCACGGCCTGCGCAACGGCAGTCCTTGGGAACCAACTTGGCACCGCTGTCGGGTTCCTGCTGCCACCCGTGCTGGTGCCTAATACCCAAAACGATACGAACCTGCTCGCCTGCAACATTTCCACTATGTTCTACGGCACCTCTGCAGTGGCGACTCTCCTGTTCATACTGACCGCGATCGCGTTCAAGGAGAAGCCGCGCTACCCACCGTCACAAGCCCAGGCTGCCTTGCAGGATTCACCACCCGAGGAATATAGCTACAAGAAGAGCATCAGGAATTTGTTCAAGAATATCCCGTTCGTGCTGTTGCTGATTACCTACGGCATTATGACAGGGGCATTTTATAGTGTGTCAACTCTCCTCAATCAAATGATTCTCACATATTATGAAGGTGAGGAAGTGAACGCAGGCAGAATCGGCCTGACCCTTGTGGTCGCCGGCATGGTAGGTAGCATCCTGTGCGGTTTGTGGCTCGACTACACCAAGACGTATAAGCAAACGACCCTTATTGTCTACATACTGTCCTTCATCGGCATGGTCATTTTCACCTTTACTCTGGATCTGAGGTACATCATTATTGTCTTCGTGACCGGTGGCGTCCTGGGGTTCTTTATGACAGGCTATCTGCCCCTGGGGTTCGAGTTCGCCGTCGAGATTACATATCCCGAGTCAGAGGGGACATCTTCCGGGCTGCTGAACGCAAGCGCCCAAATTTTCGGTATCCTGTTTACCCTGGCCCAGGGGAAATTGACCTCTGATTACGGACCGAAAGCTGGTAATATCTTCCTTTGCGTGTGGATGTTCATTGGGATTATCCTGACGGCCCTGATTAAATCCGACCTTCGGCGGCATAATATCAACATTGGGATCACCAACGTCGACGTCAAGGCCATCCCCGCCGATAGCCCGACTGATCAGGAGCCTAAGACTGTGATGCTGAGTAAACAATCTGAGAGTGCTATCTAG

## Q471A hFLVCR1

**Forward primer 1 (**5’-3’)

GTCCGAAGCGCGCTAGC

**Reverse primer 1 (**5’-3’)

CCGAAAATGGCGGCGCTTGCGTTC

**Forward primer 2 (**5’-3’)

GAACGCAAGCGCCGCCATTTTCGG

**Reverse primer 2 (**5’-3’)

GGACCTTGAAACAAAACTTCCAAAGAATTCGAG

**cDNA** (Codon Optimized; 5’-3’)

ATGGCCCGCCCCGATGACGAAGAGGGAGCTGCCGTAGCTCCTGGCCATCCACTGGCCAAGGGGTATCTGCCTCTCCCACGGGGTGCCCCTGTGGGAAAAGAATCCGTCGAACTTCAAAATGGCCCTAAGGCCGGAACTTTTCCTGTTAACGGCGCGCCTCGTGATTCTCTGGCAGCTGCTAGCGGTGTATTGGGTGGACCACAAACCCCACTTGCACCCGAAGAAGAAACACAAGCCAGATTGCTGCCAGCCGGCGCGGGCGCCGAAACCCCCGGCGCCGAATCATCACCACTTCCTCTGACAGCCCTGTCTCCTAGACGATTTGTCGTCCTGCTCATATTTAGTCTTTACTCTCTCGTGAACGCATTCCAATGGATACAGTATTCAATAATCTCAAATGTGTTTGAAGGATTTTATGGCGTGACTCTGTTGCATATAGATTGGCTTAGCATGGTCTATATGCTCGCTTATGTTCCTCTGATATTTCCAGCTACATGGCTTCTCGATACTAGGGGTCTCCGCCTTACAGCACTCCTCGGGTCAGGTCTGAATTGTCTCGGAGCATGGATAAAATGTGGATCCGTCCAACAACACCTGTTTTGGGTGACGATGCTCGGGCAATGTCTTTGTTCAGTTGCACAAGTCTTTATTCTCGGGCTGCCATCAAGAATTGCTTCTGTCTGGTTCGGCCCTAAGGAAGTAAGCACGGCCTGCGCAACGGCAGTCCTTGGGAACCAACTTGGCACCGCTGTCGGGTTCCTGCTGCCACCCGTGCTGGTGCCTAATACCCAAAACGATACGAACCTGCTCGCCTGCAACATTTCCACTATGTTCTACGGCACCTCTGCAGTGGCGACTCTCCTGTTCATACTGACCGCGATCGCGTTCAAGGAGAAGCCGCGCTACCCACCGTCACAAGCCCAGGCTGCCTTGCAGGATTCACCACCCGAGGAATATAGCTACAAGAAGAGCATCAGGAATTTGTTCAAGAATATCCCGTTCGTGCTGTTGCTGATTACCTACGGCATTATGACAGGGGCATTTT ATAGTGTGTCAACTCTCCTCAATCAAATGATTCTCACATATTATGAAGGTGAGGAAGTGAACGCAGGCAGAATCGGCCTGACCCTTGTGGTCGCCGGCATGGTAGGTAGCATCCTGTGCGGTTTGTGGCTCGACTACACCAAGACGTATAAGCAAACGACCCTTATTGTCTACATACTGTCCTTCATCGGCATGGTCATTTTCACCTTTACTCTGGATCTGAGGTACATCATTATTGTCTTCGTGACCGGTGGCGTCCTGGGGTTCTTTATGACAGGCTATCTGCCCCTGGGGTTCGAGTTCGCCGTCG AGATTACATATCCCGAGTCAGAGGGGACATCTTCCGGGCTGCTGAACGCAAGCGCC**GCC**A TTTTCGGTATCCTGTTTACCCTGGCCCAGGGGAAATTGACCTCTGATTACGGACCGAAAGCTGGTAATATCTTCCTTTGCGTGTGGATGTTCATTGGGATTATCCTGACGGCCCTGATTAAATCCGACCTTCGGCGGCATAATATCAACATTGGGATCACCAACGTCGACGTCAAGGCCATCCCCGCCGATAGCCCGACTGATCAGGAGCCTAAGACTGTGATGCTGAGTAAACAATCTGAGAGTGCTATCTAG

## Y349A hFLVCR1

**Forward primer 1 (**5’-3’)

GTCCGAAGCGCGCTAGC

**Reverse primer 1 (**5’-3’)

GTTGACACACTGGCAAATGCCCCTG

**Forward primer 2 (**5’-3’)

CAGGGGCATTTGCCAGTGTGTCAAC

**Reverse primer 2 (**5’-3’)

GGACCTTGAAACAAAACTTCCAAAGAATTCGAG

**cDNA** (Codon Optimized; 5’-3’)

ATGGCCCGCCCCGATGACGAAGAGGGAGCTGCCGTAGCTCCTGGCCATCCACTGGCCAAGGGGTATCTGCCTCTCCCACGGGGTGCCCCTGTGGGAAAAGAATCCGTCGAACTTCAAAATGGCCCTAAGGCCGGAACTTTTCCTGTTAACGGCGCGCCTCGTGATTCTCTGGCAGCTGCTAGCGGTGTATTGGGTGGACCACAAACCCCACTTGCACCCGAAGAAGAAACACAAGCCAGATTGCTGCCAGCCGGCGCGGGCGCCGAAACCCCCGGCGCCGAATCATCACCACTTCCTCTGACAGCCCTGTCTCCTAGACGATTTGTCGTCCTGCTCATATTTAGTCTTTACTCTCTCGTGAACGCATTCCAATGGATACAGTATTCAATAATCTCAAATGTGTTTGAAGGATTTTATGGCGTGACTCTGTTGCATATAGATTGGCTTAGCATGGTCTATATGCTCGCTTATGTTCCTCTGATATTTCCAGCTACATGGCTTCTCGATACTAGGGGTCTCCGCCTTACAGCACTCCTCGGGTCAGGTCTGAATTGTCTCGGAGCATGGATAAAATGTGGATCCGTCCAACAACACCTGTTTTGGGTGACGATGCTCGGGCAATGTCTTTGTTCAGTTGCACAAGTCTTTATTCTCGGGCTGCCATCAAGAATTGCTTCTGTCTGGTTCGGCCCTAAGGAAGTAAGCACGGCCTGCGCAACGGCAGTCCTTGGGAACCAACTTGGCACCGCTGTCGGGTTCCTGCTGCCACCCGTGCTGGTGCCTAATACCCAAAACGATACGAACCTGCTCGCCTGCAACATTTCCACTATGTTCTACGGCACCTCTGCAGTGGCGACTCTCCTGTTCATACTGACCGCGATCGCGTTCAAGGAGAAGCCGCGCTACCCACCGTCACAAGCCCAGGCTGCCTTGCAGGATTCACCACCCGAGGAATATAGCTACAAGAAGAGCATCAGGAATTT GTTCAAGAATATCCCGTTCGTGCTGTTGCTGATTACCTACGGCATTATGACAGGGGCATTT**G CC**AGTGTGTCAACTCTCCTCAATCAAATGATTCTCACATATTATGAAGGTGAGGAAGTGAA CGCAGGCAGAATCGGCCTGACCCTTGTGGTCGCCGGCATGGTAGGTAGCATCCTGTGCGGTTTGTGGCTCGACTACACCAAGACGTATAAGCAAACGACCCTTATTGTCTACATACTGTCCTTCATCGGCATGGTCATTTTCACCTTTACTCTGGATCTGAGGTACATCATTATTGTCTTCGTGACCGGTGGCGTCCTGGGGTTCTTTATGACAGGCTATCTGCCCCTGGGGTTCGAGTTCGCCGTCGAGATTACATATCCCGAGTCAGAGGGGACATCTTCCGGGCTGCTGAACGCAAGCGCCCAAATTTTCGGTATCCTGTTTACCCTGGCCCAGGGGAAATTGACCTCTGATTACGGACCGAAAGCTGGTAATATCTTCCTTTGCGTGTGGATGTTCATTGGGATTATCCTGACGGCCCTGATTAAATCCGACCTTCGGCGGCATAATATCAACATTGGGATCACCAACGTCGACGTCAAGGCCATCCCCG CCGATAGCCCGACTGATCAGGAGCCTAAGACTGTGATGCTGAGTAAACAATCTGAGAGTGCTATCTAG

## Q214A hFLVCR1

**Forward primer 1 (**5’-3’) GTCCGAAGCGCGCTAGC

**Reverse primer 1 (**5’-3’) CCGAGAATAAAGACGGCTGCAACTGAAC

**Forward primer 2 (**5’-3’) GTTCAGTTGCAGCCGTCTTTATTCTCGG

**Reverse primer 2 (**5’-3’) GGACCTTGAAACAAAACTTCCAAAGAATTCGAG

**cDNA** (Codon Optimized; 5’-3’) ATGGCCCGCCCCGATGACGAAGAGGGAGCTGCCGTAGCTCCTGGCCATCCACTGGCCAAGGGGTATCTGCCTCTCCCACGGGGTGCCCCTGTGGGAAAAGAATCCGTCGAACTTCAAAATGGCCCTAAGGCCGGAACTTTTCCTGTTAACGGCGCGCCTCGTGATTCTCTGGCAGCTGCTAGCGGTGTATTGGGTGGACCACAAACCCCACTTGCACCCGAAGAAGAAACACAAGCCAGATTGCTGCCAGCCGGCGCGGGCGCCGAAACCCCCGGCGCCGAATCATCACCACTTCCTCTGACAGCCCTGTCTCCTAGACGATTTGTCGTCCTGCTCATATTTAGTCTTTACTCTCTCGTGAACGCATTCCAATGGATACAGTATTCAATAATCTCAAATGTGTTTGAAGGATTTTATGGCGTGACTCTGTTGCATATAGATTGGCTTAGCATGGTCTATATGCTCGCTTATGTTCCTCTGATATTTCCAGCTACATGGCTTCTCGATACTAGGGGTCTCCGCCTTACAGCACTCCTCGGGTCAGGTCTGAATTGTCTCGGAGCATGGATAAAATGTGGATCCGTCCAACAACACCTGTTTTGGGTGACGATGCTC GGGCAATGTCTTTGTTCAGTTGCA**GCC**GTCTTTATTCTCGGGCTGCCATCAAGAATTGCTTC TGTCTGGTTCGGCCCTAAGGAAGTAAGCACGGCCTGCGCAACGGCAGTCCTTGGGAACCAACTTGGCACCGCTGTCGGGTTCCTGCTGCCACCCGTGCTGGTGCCTAATACCCAAAACGATACGAACCTGCTCGCCTGCAACATTTCCACTATGTTCTACGGCACCTCTGCAGTGGCGACTCTCCTGTTCATACTGACCGCGATCGCGTTCAAGGAGAAGCCGCGCTACCCACCGTCACAAGCCCAGGCTGCCTTGCAGGATTCACCACCCGAGGAATATAGCTACAAGAAGAGCATCAGGAATTTGTTCAAGAATATCCCGTTCGTGCTGTTGCTGATTACCTACGGCATTATGACAGGGGCATTTTATAGTGTGTCAACTCTCCTCAATCAAATGATTCTCACATATTATGAAGGTGAGGAAGTGAACGCAGGCAGAATCGGCCTGACCCTTGTGGTCGCCGGCATGGTAGGTAGCATCCTGTGCGGTTTGTGGCTCGACTACACCAAGACGTATAAGCAAACGACCCTTATTGTCTACATACTGTCCTTCATCGGCATGGTCATTTTCACCTTTACTCTGGATCTGAGGTACATCATTATTGTCTTCGTGACCGGTGGCGTCCTGGGGTTCTTTATGACAGGCTATCTGCCCCTGGGGTTCGAGTTCGCCGTCGAGATTACATATCCCGAGTCAGAGGGGACATCTTCCGGGCTGCTGAACGCAAGCGCCCAAATTTTCGGTATCCTGTTTACCCTGGCCCAGGGGAAATTGACCTCTGATTACGGACCGAAAGCTGGTAATATCTTCCTTTGCGTGTGGATGTTCATTGGGATTATCCTGACGGCCCTGATTAAATCCGACCTTCGGCGGCATAATATCAACATTGGGATCACCAACGTCGACGTCAAGGCCATCCCCGCCGATAGCCCGACTGATCAGGAGCCTAAGACTGTGATGCTGAGTAAACAATCTGAGAGTGCTATCTAG

## N245A hFLVCR1

**Forward primer 1 (**5’-3’) GTCCGAAGCGCGCTAGC

Reverse primer 1 (5’-3’)

GTGCCAAGTTGGGCCCCAAGGACTG

**Forward primer 2 (**5’-3’) CAGTCCTTGGGGCCCAACTTGGCAC

**Reverse primer 2 (**5’-3’) GGACCTTGAAACAAAACTTCCAAAGAATTCGAG

**cDNA** (Codon Optimized; 5’-3’) ATGGCCCGCCCCGATGACGAAGAGGGAGCTGCCGTAGCTCCTGGCCATCCACTGGCCAAGGGGTATCTGCCTCTCCCACGGGGTGCCCCTGTGGGAAAAGAATCCGTCGAACTTCAAAATGGCCCTAAGGCCGGAACTTTTCCTGTTAACGGCGCGCCTCGTGATTCTCTGGCAGCTGCTAGCGGTGTATTGGGTGGACCACAAACCCCACTTGCACCCGAAGAAGAAACACAAGCCAGATTGCTGCCAGCCGGCGCGGGCGCCGAAACCCCCGGCGCCGAATCATCACCACTTCCTCTGACAGCCCTGTCTCCTAGACGATTTGTCGTCCTGCTCATATTTAGTCTTTACTCTCTCGTGAACGCATTCCAATGGATACAGTATTCAATAATCTCAAATGTGTTTGAAGGATTTTATGGCGTGACTCTGTTGCATATAGATTGGCTTAGCATGGTCTATATGCTCGCTTATGTTCCTCTGATATTTCCAGCTACATGGCTTCTCGATACTAGGGGTCTCCGCCTTACAGCACTCCTCGGGTCAGGTCTGAATTGTCTCGGAGCATGGATAAAATGTGGATCCGTCCAACAACACCTGTTTTGGGTGACGATGCTCGGGCAATGTCTTTGTTCAGTTGCACAAGTCTTTATTCTCGGGCTGCCATCAAGAATTGCTTC TGTCTGGTTCGGCCCTAAGGAAGTAAGCACGGCCTGCGCAACGGCAGTCCTTGGG**GCC**CA ACTTGGCACCGCTGTCGGGTTCCTGCTGCCACCCGTGCTGGTGCCTAATACCCAAAACGATACGAACCTGCTCGCCTGCAACATTTCCACTATGTTCTACGGCACCTCTGCAGTGGCGACTCTCCTGTTCATACTGACCGCGATCGCGTTCAAGGAGAAGCCGCGCTACCCACCGTCACAAGCCCAGGCTGCCTTGCAGGATTCACCACCCGAGGAATATAGCTACAAGAAGAGCATCAGGAATTTGTTCAAGAATATCCCGTTCGTGCTGTTGCTGATTACCTACGGCATTATGACAGGGGCATTTTATAGTGTGTCAACTCTCCTCAATCAAATGATTCTCACATATTATGAAGGTGAGGAAGTGAACGCAGGCAGAATCGGCCTGACCCTTGTGGTCGCCGGCATGGTAGGTAGCATCCTGTGCGGTTTGTGGCTCGACTACACCAAGACGTATAAGCAAACGACCCTTATTGTCTACATACTGTCCTTCATCGGCATGGTCATTTTCACCTTTACTCTGGATCTGAGGTACATCATTATTGTCTTCGTGACCGGTGGCGTCCTGGGGTTCTTTATGACAGGCTATCTGCCCCTGGGGTTCGAGTTCGCCGTCGAGATTACATATCCCGAGTCAGAGGGGACATCTTCCGGGCTGCTGAACGCAAGCGCCCAAATTTTCGGTATCCTGTTTACCCTGGCCCAGGGGAAATTGACCTCTGATTACGGACCGAAAGCTGGTAATATCTTCCTTTGCGTGTGGATGTTCATTGGGATTATCCTGACGGCCCTGATTAAATCCGACCTTCGGCGGCATAATATCAACATTGGGATCACCAACGTCGACGTCAAGGCCATCCCCGCCGATAGCCCGACTGATCAGGAGCCTAAGACTGTGATGCTGAGTAAACAATCTGAGAGTGCTATCTAG

## Y153A hFLVCR1

**Forward primer 1 (**5’-3’) GTCCGAAGCGCGCTAGC

**Reverse primer 1 (**5’-3’) CATAAGCGAGCATGGCGACCATGCTAAG

**Forward primer 2 (**5’-3’) CTTAGCATGGTCGCCATGCTCGCTTATG

**Reverse primer 2 (**5’-3’) GGACCTTGAAACAAAACTTCCAAAGAATTCGAG

**cDNA** (Codon Optimized; 5’-3’) ATGGCCCGCCCCGATGACGAAGAGGGAGCTGCCGTAGCTCCTGGCCATCCACTGGCCAAGGGGTATCTGCCTCTCCCACGGGGTGCCCCTGTGGGAAAAGAATCCGTCGAACTTCAAAATGGCCCTAAGGCCGGAACTTTTCCTGTTAACGGCGCGCCTCGTGATTCTCTGGCAGCTGCTAGC GGTGTATTGGGTGGACCACAAACCCCACTTGCACCCGAAGAAGAAACACAAGCCAGATTGCTGCCAGCCGGCGCGGGCGCCGAAACCCCCGGCGCCGAATCATCACCACTTCCTCTGACAGCCCTGTCTCCTAGACGATTTGTCGTCCTGCTCATATTTAGTCTTTACTCTCTCGTGAACGCATTCCAATGGATACAGTATTCAATAATCTCAAATGTGTTTGAAGGATTTTATGGCGTGACTCTG TTGCATATAGATTGGCTTAGCATGGTC**GCC**ATGCTCGCTTATGTTCCTCTGATATTTCCAGC TACATGGCTTCTCGATACTAGGGGTCTCCGCCTTACAGCACTCCTCGGGTCAGGTCTGAATTGTCTCGGAGCATGGATAAAATGTGGATCCGTCCAACAACACCTGTTTTGGGTGACGATGCTCGGGCAATGTCTTTGTTCAGTTGCACAAGTCTTTATTCTCGGGCTGCCATCAAGAATTGCTTCTGTCTGGTTCGGCCCTAAGGAAGTAAGCACGGCCTGCGCAACGGCAGTCCTTGGGAACCAACTTGGCACCGCTGTCGGGTTCCTGCTGCCACCCGTGCTGGTGCCTAATACCCAAAACGATACGAACCTGCTCGCCTGCAACATTTCCACTATGTTCTACGGCACCTCTGCAGTGGCGACTCTCCTGTTCATACTGACCGCGATCGCGTTCAAGGAGAAGCCGCGCTACCCACCGTCACAAGCCCAGGCTGCCTTGCAGGATTCACCACCCGAGGAATATAGCTACAAGAAGAGCATCAGGAATTTGTTCAAGAATATCCCGTTCGTGCTGTTGCTGATTACCTACGGCATTATGACAGGGGCATTTTATAGTGTGTCAACTCTCCTCAATCAAATGATTCTCACATATTATGAAGGTGAGGAAGTGAACGCAGGCAGAATCGGCCTGACCCTTGTGGTCGCCGGCATGGTAGGTAGCATCCTGTGCGGTTTGTGGCTCGACTACACCAAGACGTATAAGCAAACGACCCTTATTGTCTACATACTGTCCTTCATCGGCATGGTCATTTTCACCTTTACTCTGGATCTGAGGTACATCATTATTGTCTTCGTGACCGGTGGCGTCCTGGGGTTCTTTATGACAGGCTATCTGCCCCTGGGGTTCGAGTTCGCCGTCGAGATTACATATCCCGAGTCAGAGGGGACATCTTCCGGGCTGCTGAACGCAAGCGCCCAAATTTTCGGTATCCTGTTTACCCTGGCCCAGGGGAAATTGACCTCTGATTACGGACCGAAAGCTGGTAATATCTTCCTTTGCGTGTGGATGTTCATTGGGATTATCCTGACGGCCCTGATTAAATCCGACCTTCGGCGGCATAATATCAACATTGGGATCACCAACGTCGACGTCAAGGCCATCCCCGCCGATAGCCCGACTGATCAGGAGCCTAAGACTGTGATGCTGAGTAAACAATCTGAGAGTGCTATCTAG

## N121D hFLVCR1

**Forward primer 1 (**5’-3’) GTCCGAAGCGCGCTAGC

**Reverse primer 1 (**5’-3’) CATTGGAATGCGTCCACGAGAGAG

**Forward primer 2 (**5’-3’) CTCTCTCGTGGACGCATTCCAATG

**Reverse primer 2 (**5’-3’) GGACCTTGAAACAAAACTTCCAAAGAATTCGAG

**cDNA** (Codon Optimized; 5’-3’) ATGGCCCGCCCCGATGACGAAGAGGGAGCTGCCGTAGCTCCTGGCCATCCACTGGCCAAGGGGTATCTGCCTCTCCCACGGGGTGCCCCTGTGGGAAAAGAATCCGTCGAACTTCAAAATGGCCCTAAGGCCGGAACTTTTCCTGTTAACGGCGCGCCTCGTGATTCTCTGGCAGCTGCTAGCGGTGTATTGGGTGGACCACAAACCCCACTTGCACCCGAAGAAGAAACACAAGCCAGATTGCTGCCAGCCGGCGCGGGCGCCGAAACCCCCGGCGCCGAATCATCACCACTTCCTCTGACAG CCCTGTCTCCTAGACGATTTGTCGTCCTGCTCATATTTAGTCTTTACTCTCTCGTG**GAC**GCAT TCCAATGGATACAGTATTCAATAATCTCAAATGTGTTTGAAGGATTTTATGGCGTGACTCTGTTGCATATAGATTGGCTTAGCATGGTCTATATGCTCGCTTATGTTCCTCTGATATTTCCAGCTACATGGCTTCTCGATACTAGGGGTCTCCGCCTTACAGCACTCCTCGGGTCAGGTCTGAATTGTCTCGGAGCATGGATAAAATGTGGATCCGTCCAACAACACCTGTTTTGGGTGACGATGCTCGGGCAATGTCTTTGTTCAGTTGCACAAGTCTTTATTCTCGGGCTGCCATCAAGAATTGCTTCTGTCTGGTTCGGCCCTAAGGAAGTAAGCACGGCCTGCGCAACGGCAGTCCTTGGGAACCAA CTTGGCACCGCTGTCGGGTTCCTGCTGCCACCCGTGCTGGTGCCTAATACCCAAAACGATACGAACCTGCTCGCCTGCAACATTTCCACTATGTTCTACGGCACCTCTGCAGTGGCGACTCTCCTGTTCATACTGACCGCGATCGCGTTCAAGGAGAAGCCGCGCTACCCACCGTCACAAGCCCAGGCTGCCTTGCAGGATTCACCACCCGAGGAATATAGCTACAAGAAGAGCATCAGGAATTTGTTCAAGAATATCCCGTTCGTGCTGTTGCTGATTACCTACGGCATTATGACAGGGGCATTTTATAGTGTGTCAACTCTCCTCAATCAAATGATTCTCACATATTATGAAGGTGAGGAAGTGAACGCAGGCAGAATCGGCCTGACCCTTGTGGTCGCCGGCATGGTAGGTAGCATCCTGTGCGGTTTGTGGCTCGACTACACCAAGACGTATAAGCAAACGACCCTTATTGTCTACATACTGTCCTTCATCGGCATGGTCATTTTCACCTTTACTCTGGATCTGAGGTACATCATTATTGTCTTCGTGACCGGTGGCGTCCTGGGGTTCTTTATGACAGGCTATCTGCCCCTGGGGTTCGAGTTCGCCGTCGAGATTACATATCCCGAGTCAGAGGGGACATCTTCCGGGCTGCTGAACGCAAGCGCCCAAATTTTCGGTATCCTGTTTACCCTGGCCCAGGGGAAATTGACCTCTGATTACGGACCGAAAGCTGGTAATATCTTCCTTTGCGTGTGGATGTTCATTGGGATTATCCTGACGGCCCTGATTAAATCCGACCTTCGGCGGCATAATATCAACATTGGGATCACCAACGTCGACGTCAAGGCCATCCCCGCCGATAGCCCGACTGATCAGGAGCCTAAGACTGTGATGCTGAGTAAACAATCTGAGAGTGCTATCTAG

## C192R hFLVCR1

**Forward primer 1 (**5’-3’) GTCCGAAGCGCGCTAGC

**Reverse primer 1 (**5’-3’) GGACGGATCCCCGTTTTATCCATG

**Forward primer 2 (**5’-3’) catggataaaaCGGggatccgtcc **Reverse primer 2 (**5’-3’)

CATGGATAAAACGGGGATCCGTCC

**cDNA** (Codon Optimized; 5’-3’) ATGGCCCGCCCCGATGACGAAGAGGGAGCTGCCGTAGCTCCTGGCCATCCACTGGCCAAGGGGTATCTGCCTCTCCCACGGGGTGCCCCTGTGGGAAAAGAATCCGTCGAACTTCAAAATGGCCCTAAGGCCGGAACTTTTCCTGTTAACGGCGCGCCTCGTGATTCTCTGGCAGCTGCTAGCGGTGTATTGGGTGGACCACAAACCCCACTTGCACCCGAAGAAGAAACACAAGCCAGATTGCTGCCAGCCGGCGCGGGCGCCGAAACCCCCGGCGCCGAATCATCACCACTTCCTCTGACAGCCCTGTCTCCTAGACGATTTGTCGTCCTGCTCATATTTAGTCTTTACTCTCTCGTGAACGCATTCCAATGGATACAGTATTCAATAATCTCAAATGTGTTTGAAGGATTTTATGGCGTGACTCTGTTGCATATAGATTGGCTTAGCATGGTCTATATGCTCGCTTATGTTCCTCTGATATTTCCAGCTACATGGCTTCTCGATACTAGGGGTCTCCGCCTTACAGCACTCCTCGGGTCAGGTCTGAATTG TCTCGGAGCATGGATAAAA**CGG**GGATCCGTCCAACAACACCTGTTTTGGGTGACGATGCTC GGGCAATGTCTTTGTTCAGTTGCACAAGTCTTTATTCTCGGGCTGCCATCAAGAATTGCTTCTGTCTGGTTCGGCCCTAAGGAAGTAAGCACGGCCTGCGCAACGGCAGTCCTTGGGAACCAACTTGGCACCGCTGTCGGGTTCCTGCTGCCACCCGTGCTGGTGCCTAATACCCAAAACGATACGAACCTGCTCGCCTGCAACATTTCCACTATGTTCTACGGCACCTCTGCAGTGGCGACTCTCCTGTTCATACTGACCGCGATCGCGTTCAAGGAGAAGCCGCGCTACCCACCGTCACAAGCCCAGGCTGCCTTGCAGGATTCACCACCCGAGGAATATAGCTACAAGAAGAGCATCAGGAATTTGTTCAAGAATATCCCGTTCGTGCTGTTGCTGATTACCTACGGCATTATGACAGGGGCATTTTATAGTGTGTCAACTCTCCTCAATCAAATGATTCTCACATATTATGAAGGTGAGGAAGTGAACGCAGGCAGAATCGGCCTGACCCTTGTGGTCGCCGGCATGGTAGGTAGCATCCTGTGCGGTTTGTGGCTCGACTACACCAAGACGTATAAGCAAACGACCCTTATTGTCTACATACTGTCCTTCATCGGCATGGTCATTTTCACCTTTACTCTGGATCTGAGGTACATCATTATTGTCTTCGTGAC CGGTGGCGTCCTGGGGTTCTTTATGACAGGCTATCTGCCCCTGGGGTTCGAGTTCGCCGTCGAGATTACATATCCCGAGTCAGAGGGGACATCTTCCGGGCTGCTGAACGCAAGCGCCCAAATTTTCGGTATCCTGTTTACCCTGGCCCAGGGGAAATTGACCTCTGATTACGGACCGAAAGCTGGTAATATCTTCCTTTGCGTGTGGATGTTCATTGGGATTATCCTGACGGCCCTGATTAAATCCGACCTTCGGCGGCATAATATCAACATTGGGATCACCAACGTCGACGTCAAGGCCATCCCCGCCGATAGCCCGACTGATCAGGAGCCTAAGACTGTGATGCTGAGTAAACAATCTGAGAGTGCTATCTAG

## N467A hFLVCR1

**Forward primer 1 (**5’-3’) GTCCGAAGCGCGCTAGC

**Reverse primer 1 (**5’-3’) GGGCGCTTGCGGCCAGCAGCCCGG

**Forward primer 2 (**5’-3’) CCGGGCTGCTGGCCGCAAGCGCCC

**Reverse primer 2 (**5’-3’) GGACCTTGAAACAAAACTTCCAAAGAATTCGAG

**cDNA** (Codon Optimized; 5’-3’) ATGGCCCGCCCCGATGACGAAGAGGGAGCTGCCGTAGCTCCTGGCCATCCACTGGCCAAGGGGTATCTGCCTCTCCCACGGGGTGCCCCTGTGGGAAAAGAATCCGTCGAACTTCAAAATGGCCCTAAGGCCGGAACTTTTCCTGTTAACGGCGCGCCTCGTGATTCTCTGGCAGCTGCTAGCGGTGTATTGGGTGGACCACAAACCCCACTTGCACCCGAAGAAGAAACACAAGCCAGATTGCTGCCAGCCGGCGCGGGCGCCGAAACCCCCGGCGCCGAATCATCACCACTTCCTCTGACAGCCCTGTCTCCTAGACGATTTGTCGTCCTGCTCATATTTAGTCTTTACTCTCTCGTGAACGCATTCCAATGGATACAGTATTCAATAATCTCAAATGTGTTTGAAGGATTTTATGGCGTGACTCTGTTGCATATAGATTGGCTTAGCATGGTCTATATGCTCGCTTATGTTCCTCTGATATTTCCAGCTACATGGCTTCTCGATACTAGGGGTCTCCGCCTTACAGCACTCCTCGGGTCAGGTCTGAATTGTCTCGGAGCATGGATAAAATGTGGATCCGTCCAACAACACCTGTTTTGGGTGACGATGCTCGGGCAATGTCTTTGTTCAGTTGCACAAGTCTTTATTCTCGGGCTGCCATCAAGAATTGCTTCTGTCTGGTTCGGCCCTAAGGAAGTAAGCACGGCCTGCGCAACGGCAGTCCTTGGGAACCAACTTGGCACCGCTGTCGGGTTCCTGCTGCCACCCGTGCTGGTGCCTAATACCCAAAACGATACGAACCTGCTCGCCTGCAACATTTCCACTATGTTCTACGGCACCTCTGCAGTGGCGACTCTCCTGTTCATACTGACCGCGATCGCGTTCAAGGAGAAGCCGCGCTACCCACCGTCACAAGCCCAGGCTGCCTTGCAGGATTCACCACCCGAGGAATATAGCTACAAGAAGAGCATCAGGAATTTGTTCAAGAATATCCCGTTCGTGCTGTTGCTGATTACCTACGGCATTATGACAGGGGCATTTTATAGTGTGTCAACTCTCCTCAATCAAATGATTCTCACATATTATGAAGGTGAGGAAGTGAACGCAGGCAGAATCGGCCTGACCCTTGTGGTCGCCGGCATGGTAGGTAGCATCCTGTGCGGTTTGTGGCTCGACTACACCAAGACGTATAAGCAAACGACCCTTATTGTCTACATACTGTCCTTCATCGGCATGGTCATTTTCACCTTTACTCTGGATCTGAGGTACATCATTATTGTCTTCGTGACCGGTGGCGTCCTGGGGTTCTTTATGACAGGCTATCTGCCCCTGGGGTTCGAGTTCGCCGTCG AGATTACATATCCCGAGTCAGAGGGGACATCTTCCGGGCTGCTG**GCC**GCAAGCGCCCAAA TTTTCGGTATCCTGTTTACCCTGGCCCAGGGGAAATTGACCTCTGATTACGGACCGAAAGCTGGTAATATCTTCCTTTGCGTGTGGATGTTCATTGGGATTATCCTGACGGCCCTGATTAAATCCGACCTTCGGCGGCATAATATCAACATTGGGATCACCAACGTCGACGTCAAGGCCATCCCCGCCGATAGCCCGACTGATCAGGAGCCTAAGACTGTGATGCTGAGTAAACAATCTGAGAGTGCTATCTAG

## A241T hFLVCR1

**Forward primer 1** (5’-3’) GTCCGAAGCGCGCTAGC

**Reverse primer 1** (5’-3’) CCCAAGGACGGTCGTTGCGCAG

**Forward primer 2** (5’-3’) CTGCGCAACGACCGTCCTTGGG

**Reverse primer 2** (5’-3’) GGACCTTGAAACAAAACTTCCAAAGAATTCGAG

**cDNA** (Codon Optimized; 5’-3’) ATGGCCCGCCCCGATGACGAAGAGGGAGCTGCCGTAGCTCCTGGCCATCCACTGGCCAAGGGGTATCTGCCTCTCCCACGGGGTGCCCCTGTGGGAAAAGAATCCGTCGAACTTCAAAATGGCCCTAAGGCCGGAACTTTTCCTGTTAACGGCGCGCCTCGTGATTCTCTGGCAGCTGCTAGCGGTGTATTGGGTGGACCACAAACCCCACTTGCACCCGAAGAAGAAACACAAGCCAGATTGCTGCCAGCCGGCGCGGGCGCCGAAACCCCCGGCGCCGAATCATCACCACTTCCTCTGACAGCCCTGTCTCCTAGACGATTTGTCGTCCTGCTCATATTTAGTCTTTACTCTCTCGTGAACGCATTCCAATGGATACAGTATTCAATAATCTCAAATGTGTTTGAAGGATTTTATGGCGTGACTCTGTTGCATATAGATTGGCTTAGCATGGTCTATATGCTCGCTTATGTTCCTCTGATATTTCCAGCTACATGGCTTCTCGATACTAGGGGTCTCCGCCTTACAGCACTCCTCGGGTCAGGTCTGAATTGTCTCGGAGCATGGATAAAATGTGGATCCGTCCAACAACACCTGTTTTGGGTGACGATGCTCGGGCAATGTCTTTGTTCAGTTGCACAAGTCTTTATTCTCGGGCTGCCATCAAGAATTGCTTC TGTCTGGTTCGGCCCTAAGGAAGTAAGCACGGCCTGCGCAACG**ACC**GTCCTTGGGAACCAA CTTGGCACCGCTGTCGGGTTCCTGCTGCCACCCGTGCTGGTGCCTAATACCCAAAACGATACGAACCTGCTCGCCTGCAACATTTCCACTATGTTCTACGGCACCTCTGCAGTGGCGACTCTCCTGTTCATACTGACCGCGATCGCGTTCAAGGAGAAGCCGCGCTACCCACCGTCACAAGCCCAGGCTGCCTTGCAGGATTCACCACCCGAGGAATATAGCTACAAGAAGAGCATCAGGAATTTGTTCAAGAATATCCCGTTCGTGCTGTTGCTGATTACCTACGGCATTATGACAGGGGCATTTTATAGTGTGTCAACTCTCCTCAATCAAATGATTCTCACATATTATGAAGGTGAGGAAGTGAACGCAGGCAGAATCGGCCTGACCCTTGTGGTCGCCGGCATGGTAGGTAGCATCCTGTGCGGTTTGTGGCTCGACTACACCAAGACGTATAAGCAAACGACCCTTATTGTCTACATACTGTCCTTCATCGGCATGGTCATTTTCACCTTTACTCTGGATCTGAGGTACATCATTATTGTCTTCGTGACCGGTGGCGTCCTGGGGTTCTTTATGACAGGCTATCTGCCCCTGGGGTTCGAGTTCGCCGTCGAGATTACATATCCCGAGTCAGAGGGGACATCTTCCGGGCTGCTGAACGCAAGCGCCCAAATTTTCGGTATCCTGTTTACCCTGGCCCAGGGGAAATTGACCTCTGATTACGGACCGAAAGCTGGTAATATCTTCCTTTGCGTGTGGATGTTCATTGGGATTATCCTGACGGCCCTGATTAAATCCGACCTTCGGCGGCATAATATCAACATTGGGATCACCAACGTCGACGTCAAGGCCATCCCCGCCGATAGCCCGACTGATCAGGAGCCTAAGACTGTGATGCTGAGTAAACAATCTGAGAGTGCTATCTAG

## DATA AVAILIBILITY

Cryo-EM maps have been deposited in the EMDB under accession codes EMD-42107 (choline-bound FLVCR1), EMD-42108 (ethanolamine-bound FLVCR1), EMD-42109 (endogenous choline-bound FLVCR1), EMD-42110 (endogenous choline-bound FLVCR1, from images collected in 1 mM ethanolamine) and EMD-42111 (endogenous ligand-bound FLVCR1). Atomic coordinates have been deposited in the PDB under accession codes 8UBW (choline-bound FLVCR1), 8UBX (ethanolamine-bound FLVCR1), 8UBY (endogenous choline-bound FLVCR1), 8UBZ (endogenous choline-bound FLVCR1, from images collected in 1 mM ethanolamine) and 8UC0 (endogenous ligand-bound FLVCR1). Source data is available with this paper.

## CODE AVAILIBILITY

The code used for the Github link to the code for FLVCR1 DepMap Coessentiality analysis is available at https://github.com/artemkhan/Coessentiality_DepMAP_FLVCR1.git.

## ACKNOWLEDGEMENTS

We thank M. J. de la Cruz and the SEMC staff for help with data acquisition; the Memorial Sloan Kettering Cancer Center HPC group for assistance with data processing and the members of the laboratories for comments on the manuscript and L. Finley for helpful discussions. R.H. is supported by the NIH-National Cancer Institute Cancer Center Support Grant P30-CA008748 and is a Searle Scholar. T.K. is supported by NIH/NIDDK (F32 DK127836), the Shapiro-Silverberg Fund for the Advancement of Translational Research and a Merck Postdoctoral Fellowship at The Rockefeller University. K.B. is supported by the NIH/NIDDK (R01 DK123323-01), Mark Foundation Emerging Leader Award; and is a Searle and Pew-Stewart Scholar. Some of this work was performed at the Simons Electron Microscopy Center at the New York Structural Biology Center, with major support from the Simons Foundation (SF349247).

## AUTHOR CONTRIBUTIONS

Conceptualization, Y.S. and R.H.; Methodology Y.S., T.K., A.K., K.B. and R.H., Formal Analysis, Y.S. and R.H.; Investigation, Y.S., T.K., K.B. and R.H.; Writing – Original Draft, Y.S. and R.H.; Funding Acquisition, T.K., K.B. and R.H.

## Extended Data

**Extended Data Table 1.**
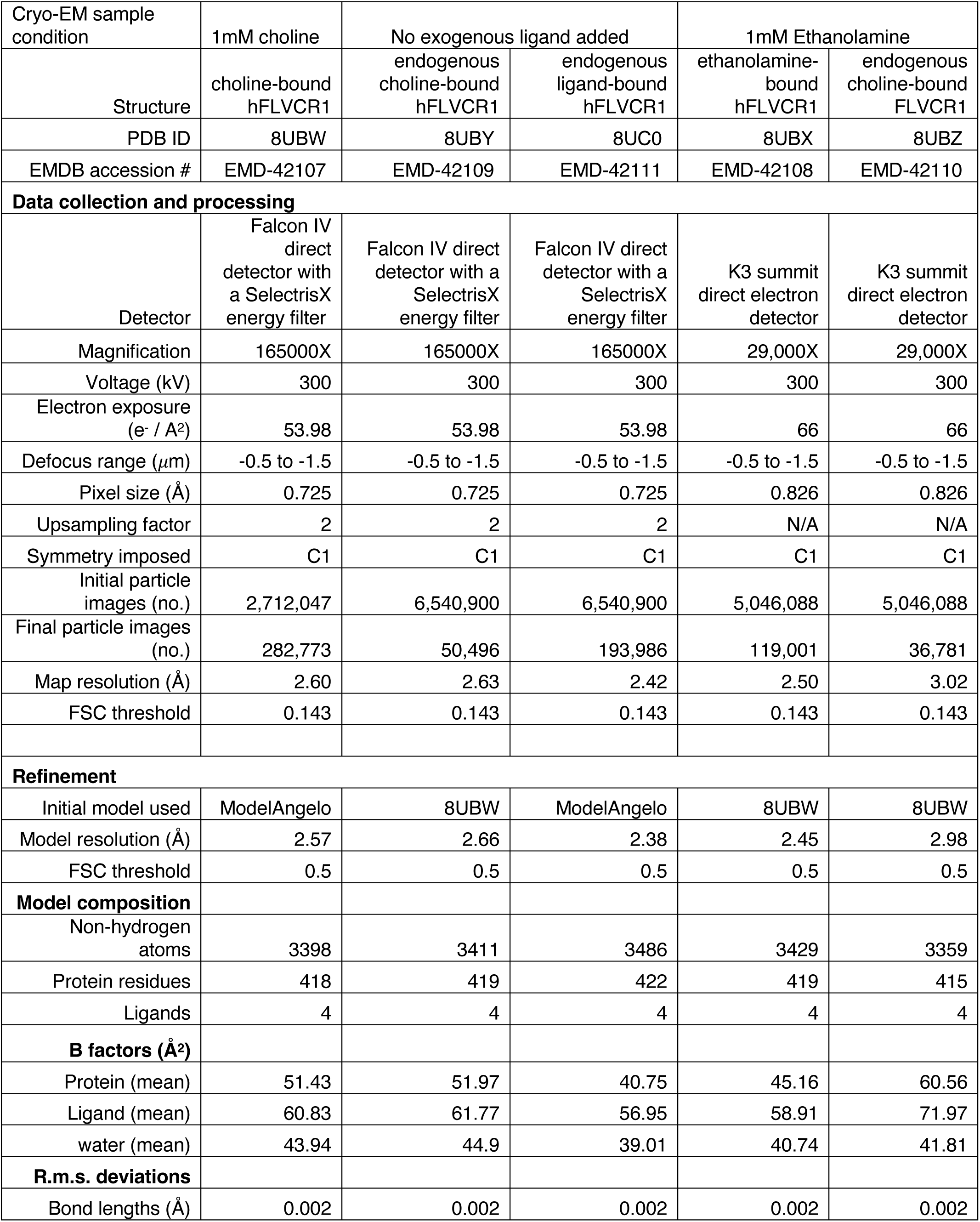
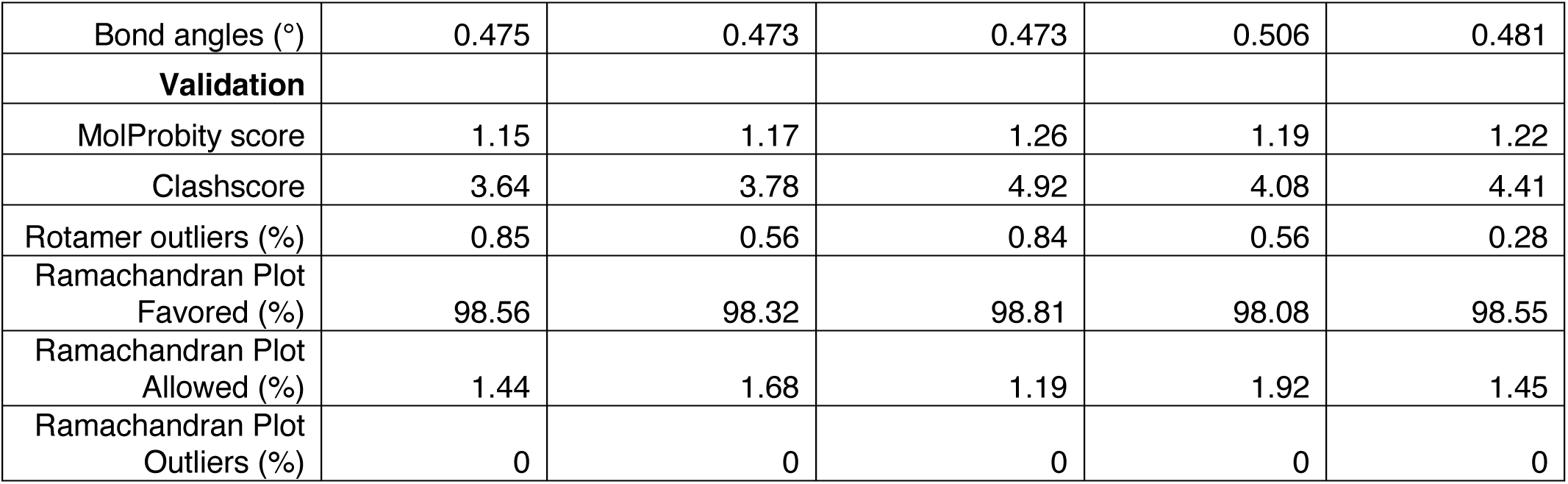
Cryo-EM data collection, refinement, and validation statistics.

| Cryo-EM sample condition | 1mM choline | No exogenous ligand added |  | 1mM Ethanolamine |  |
| --- | --- | --- | --- | --- | --- |
| Structure | choline-bound hFLVCR1 | endogenous choline-bound hFLVCR1 | endogenous ligand-bound hFLVCR1 | ethanolamine-bound hFLVCR1 | endogenous choline-bound FLVCR1 |
| PDB ID | 8UBW | 8UBY | 8UC0 | 8UBX | 8UBZ |
| EMDB accession # | EMD-42107 | EMD-42109 | EMD-42111 | EMD-42108 | EMD-42110 |
| Data collection and processing |  |  |  |  |  |
| Detector | Falcon IV direct detector with a SelectrisX energy filter | Falcon IV direct detector with a SelectrisX energy filter | Falcon IV direct detector with a SelectrisX energy filter | K3 summit direct electron detector | K3 summit direct electron detector |
| Magnification | 165000X | 165000X | 165000X | 29,000X | 29,000X |
| Voltage (kV) | 300 | 300 | 300 | 300 | 300 |
| Electron exposure (e<sup>-</sup> / Å<sup>2</sup>) | 53.98 | 53.98 | 53.98 | 66 | 66 |
| Defocus range (μm) | -0.5 to -1.5 | -0.5 to -1.5 | -0.5 to -1.5 | -0.5 to -1.5 | -0.5 to -1.5 |
| Pixel size (Å) | 0.725 | 0.725 | 0.725 | 0.826 | 0.826 |
| Upsampling factor | 2 | 2 | 2 | N/A | N/A |
| Symmetry imposed | C1 | C1 | C1 | C1 | C1 |
| Initial particle images (no.) | 2,712,047 | 6,540,900 | 6,540,900 | 5,046,088 | 5,046,088 |
| Final particle images (no.) | 282,773 | 50,496 | 193,986 | 119,001 | 36,781 |
| Map resolution (Å) | 2.60 | 2.63 | 2.42 | 2.50 | 3.02 |
| FSC threshold | 0.143 | 0.143 | 0.143 | 0.143 | 0.143 |
| Refinement |  |  |  |  |  |
| Initial model used | ModelAngelo | 8UBW | ModelAngelo | 8UBW | 8UBW |
| Model resolution (Å) | 2.57 | 2.66 | 2.38 | 2.45 | 2.98 |
| FSC threshold | 0.5 | 0.5 | 0.5 | 0.5 | 0.5 |
| Model composition |  |  |  |  |  |
| Non-hydrogen atoms | 3398 | 3411 | 3486 | 3429 | 3359 |
| Protein residues | 418 | 419 | 422 | 419 | 415 |
| Ligands | 4 | 4 | 4 | 4 | 4 |
| B factors (Å<sup>2</sup>) |  |  |  |  |  |
| Protein (mean) | 51.43 | 51.97 | 40.75 | 45.16 | 60.56 |
| Ligand (mean) | 60.83 | 61.77 | 56.95 | 58.91 | 71.97 |
| water (mean) | 43.94 | 44.9 | 39.01 | 40.74 | 41.81 |
| R.m.s. deviations |  |  |  |  |  |
| Bond lengths (Å) | 0.002 | 0.002 | 0.002 | 0.002 | 0.002 |
| Bond angles (°) | 0.475 | 0.473 | 0.473 | 0.506 | 0.481 |
| **Validation** |  |  |  |  |  |
| MolProbity score | 1.15 | 1.17 | 1.26 | 1.19 | 1.22 |
| Clashscore | 3.64 | 3.78 | 4.92 | 4.08 | 4.41 |
| Rotamer outliers (%) | 0.85 | 0.56 | 0.84 | 0.56 | 0.28 |
| Ramachandran Plot<br>Favored (%) | 98.56 | 98.32 | 98.81 | 98.08 | 98.55 |
| Ramachandran Plot<br>Allowed (%) | 1.44 | 1.68 | 1.19 | 1.92 | 1.45 |
| Ramachandran Plot<br>Outliers (%) | 0 | 0 | 0 | 0 | 0 |

**Extended Data Fig. 1.**
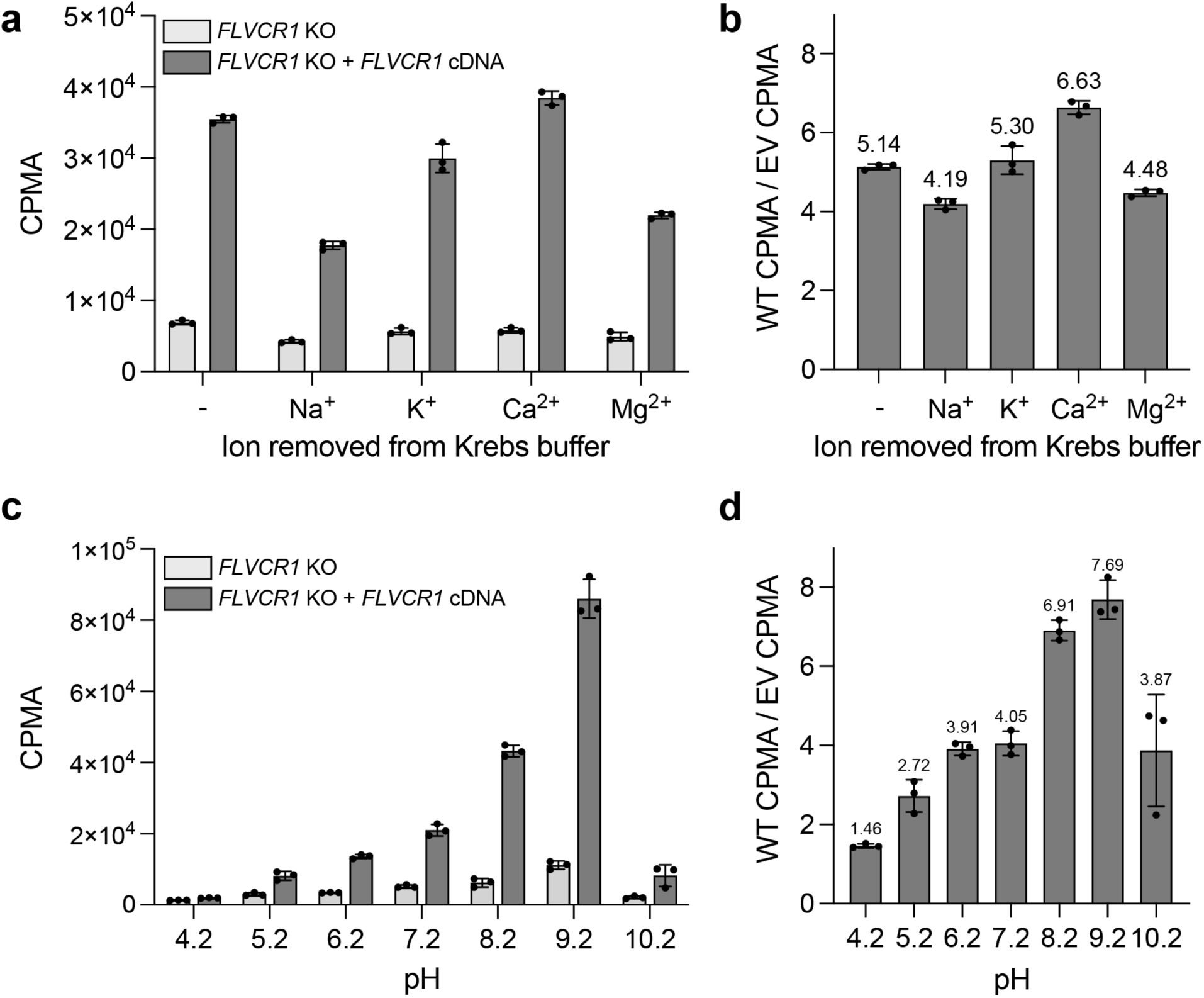
Regulation of FLVCR1-mediated choline uptake. **a**, Raw uptake of 21.5 μM choline by *FLVCR1*-knockout HEK293T cells expressing a vector control (white) or *FLVCR1* cDNA (grey) after 30 minutes in Krebs buffer or Krebs buffer lacking Na^+^, K^+^, Ca^2+^, or Mg^2+^. Data shown as mean ± standard deviation; n=3. **b**, Ratio of uptake of 21.5 μM choline by *FLVCR1*-knockout HEK293T cells expressing *FLVCR1* cDNA to the average uptake of 21.5 μM choline by *FLVCR1*-knockout HEK293T cells expressing a vector control after 30 minutes in Krebs buffer or Krebs buffer lacking Na^+^, K^+^, Ca^2+^ or Mg^2+^. Data shown as mean ± standard deviation; n=3. **c**, Raw uptake of 21.5 μM choline by *FLVCR1*-knockout HEK293T cells expressing a vector control (white) or *FLVCR1* cDNA (grey) after 30 minutes in Krebs buffer at pH 4.2 – 10.2. Data shown as mean ± standard deviation; n=3. **d**, Ratio of uptake of 21.5 μM choline by *FLVCR1*-knockout HEK293T cells expressing *FLVCR1* cDNA to the average uptake of 21.5 μM choline by *FLVCR1*-knockout HEK293T cells expressing a vector control after 30 minutes Krebs buffer at pH 4.2 – 10.2. Data shown as mean ± standard deviation; n=3.

**Extended Data Fig. 2.**
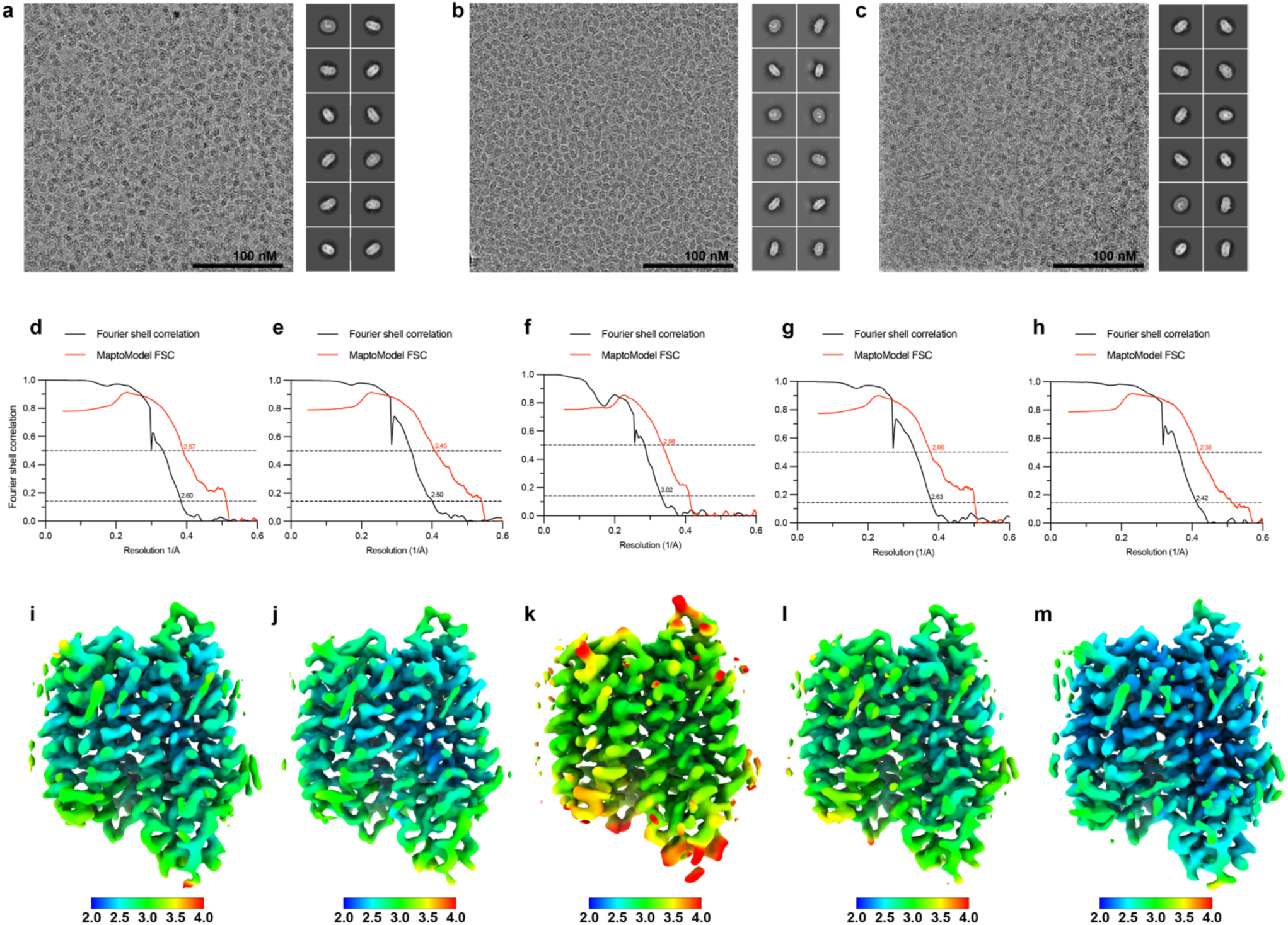
Validation of cryo-EM structures of FLVCR1. **a-c** Representative cryo-EM images and two-dimensional class averages of human FLVCR1 in 1 mM choline (a), 1 mM ethanolamine (b) or without added substrate (c). **d-h**, Plots showing Fourier shell correlations between two independent half-maps (black) and between density-modified map and refined atomic model (red) for choline-bound FLVCR1 (d), ethanolamine-bound FLVCR1 (e), endogenous choline-bound FLVCR1 obtained from FLVCR1 incubated with 1mM ethanolamine (f), endogenous choline-bound FLVCR1 obtained from FLVCR1 without exogenous ligand incubation (g), and endogenous ligand-bound FLVCR1 obtained from FLVCR1 without exogenous ligand incubation (h). Dashed lines are indicated at FSC = 0.5 and FSC = 0.143. **i-m**, Local resolution plots of choline-bound FLVCR1 (i), ethanolamine-bound FLVCR1 (j), endogenous-choline bound FLVCR1 obtained from FLVCR1 incubated with 1mM ethanolamine (k), endogenous choline-bound FLVCR1 obtained from FLVCR1 without exogenous substrate incubation (l), and endogenous ligand-bound FLVCR1 obtained from FLVCR1 without exogenous ligand incubation (m).

**Extended Data Fig. 3.**
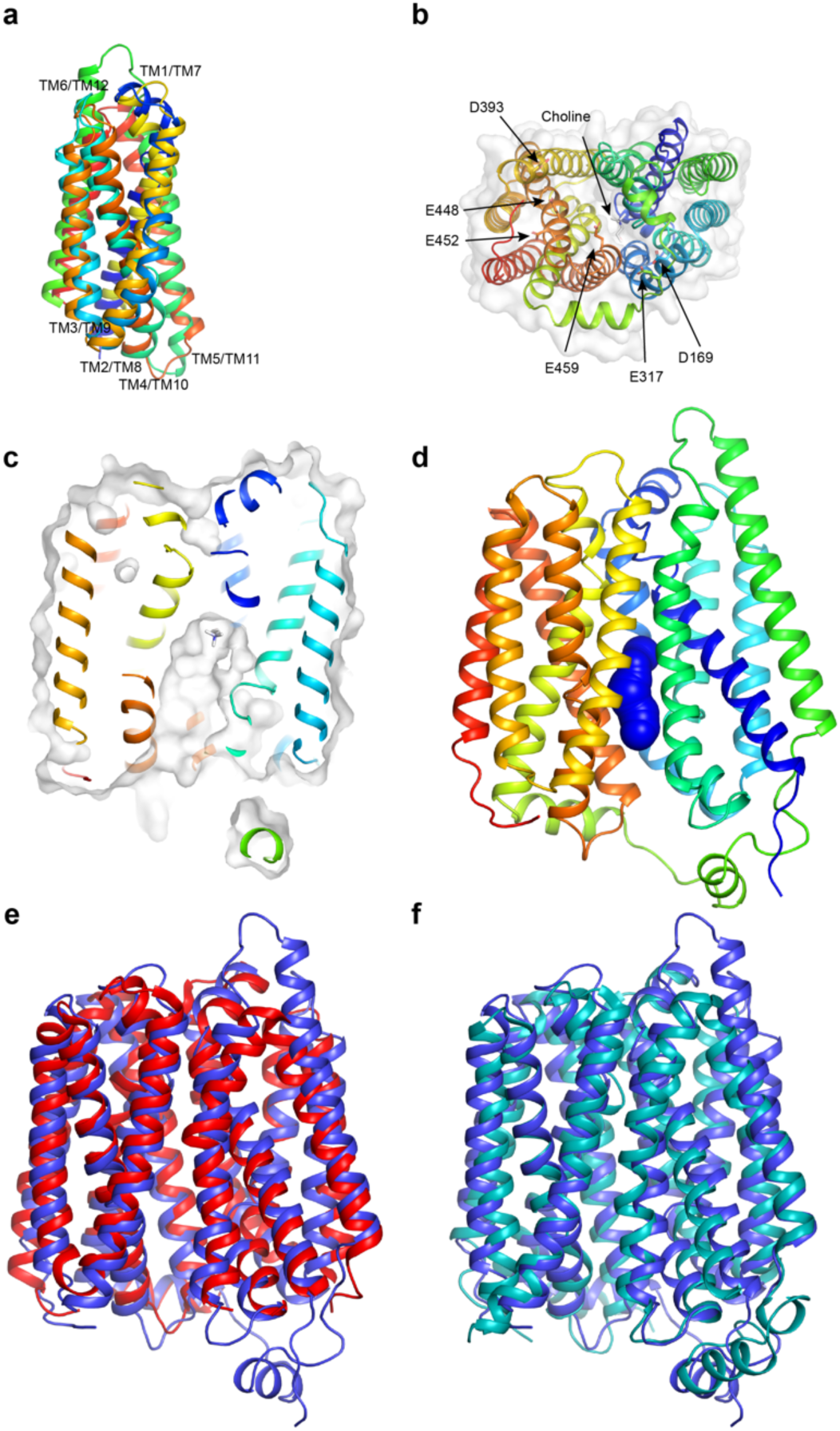
Choline-bound FLVCR1 adopts an inward-facing conformation. **a**, Superposition of TM1-6 and TM7-12. RMSD = 3.5 Å. **b**, Central cavity viewed from the cytosolic side. Aspartate and glutamate residues in and near the entrance to the central cavity are shown as sticks. **c**, Central section of choline-bound FLVCR1 is contiguous with choline shown as sticks. **d**, Blue spheres depict the minimum radius of the central cavity in the choline-bound state as a function of position. **e-f**, Superposition of choline-bound FLVCR1 (blue) with *E. coli* SotB (D; PDB: 6KKL; RMSD = 2.3 Å; red) (e) or *E. coli* DgoT (E; PDB: 6E9N; RMSD = 2.6 Å; cyan) (f).

**Extended Data Fig. 4.**
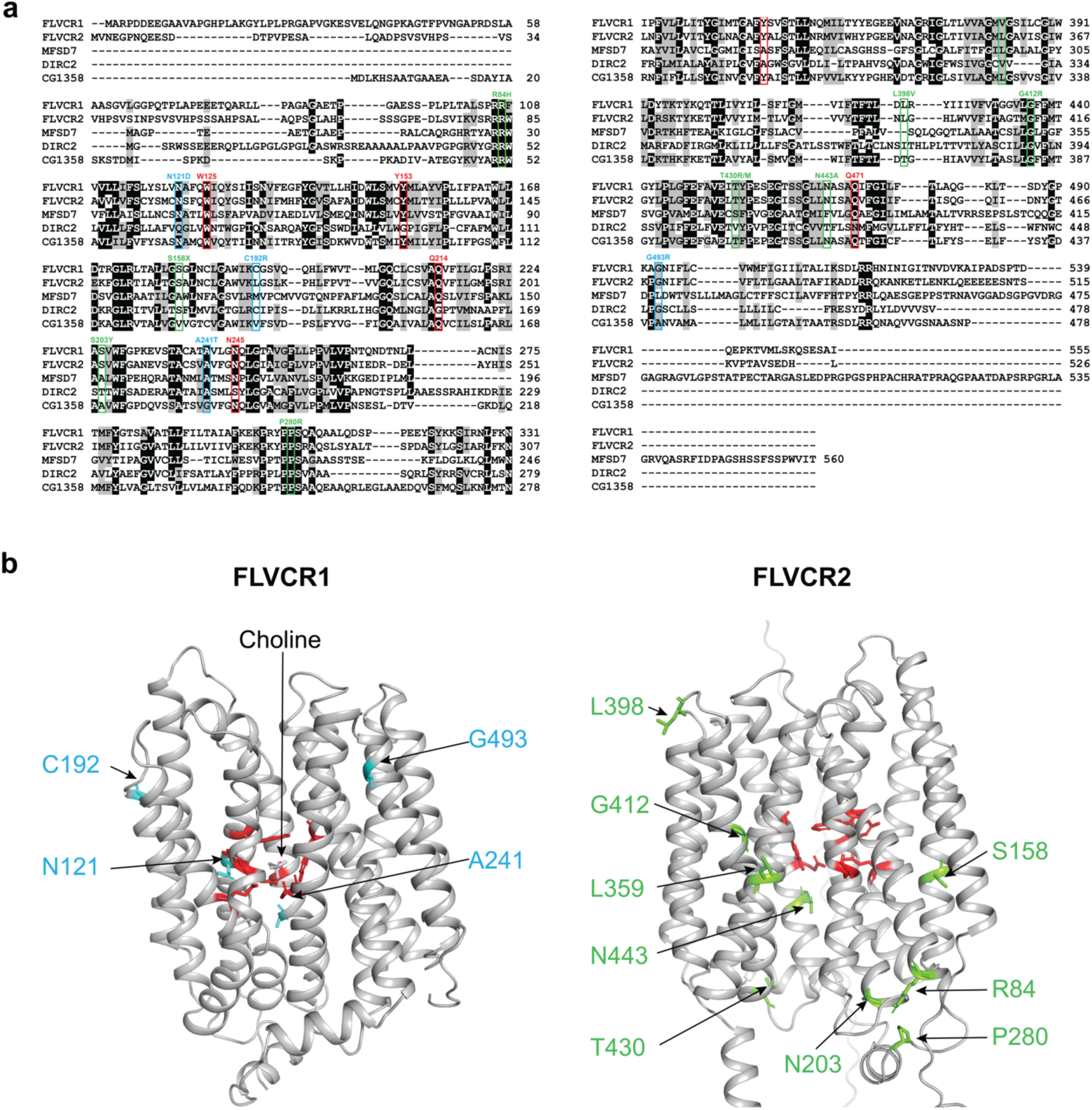
Sequence alignment of SLC49A family members and identified disease-associated mutations. **a**, Sequence alignment of human FLVCR1 (SLC49A1), human FLVCR2 (SLC49A2), human MFSD7 (SLC49A3), human DIRC2 (SLC49A4) and *Drosophila* CG1358 performed using Clustal Omega ^49^ and pyBoxshade (https://github.com/mdbaron42/pyBoxshade). Substrate-binding site residues are highlighted by red boxes. Residues whose mutation leads to PCARP or Fowler syndrome are highlighted by blue and green boxes, respectively. **b**, Structure of choline-bound FLVCR1 (left) and Alphafold2 model of FLVCR2 (right) ^50^ with residues whose mutation leads to PCARP and Fowler syndrome highlighted in blue and green, respectively. FLVCR1 substrate binding site residues and the corresponding residues in FLVCR2 are highlighted in red.

**Extended Data Fig. 5.**
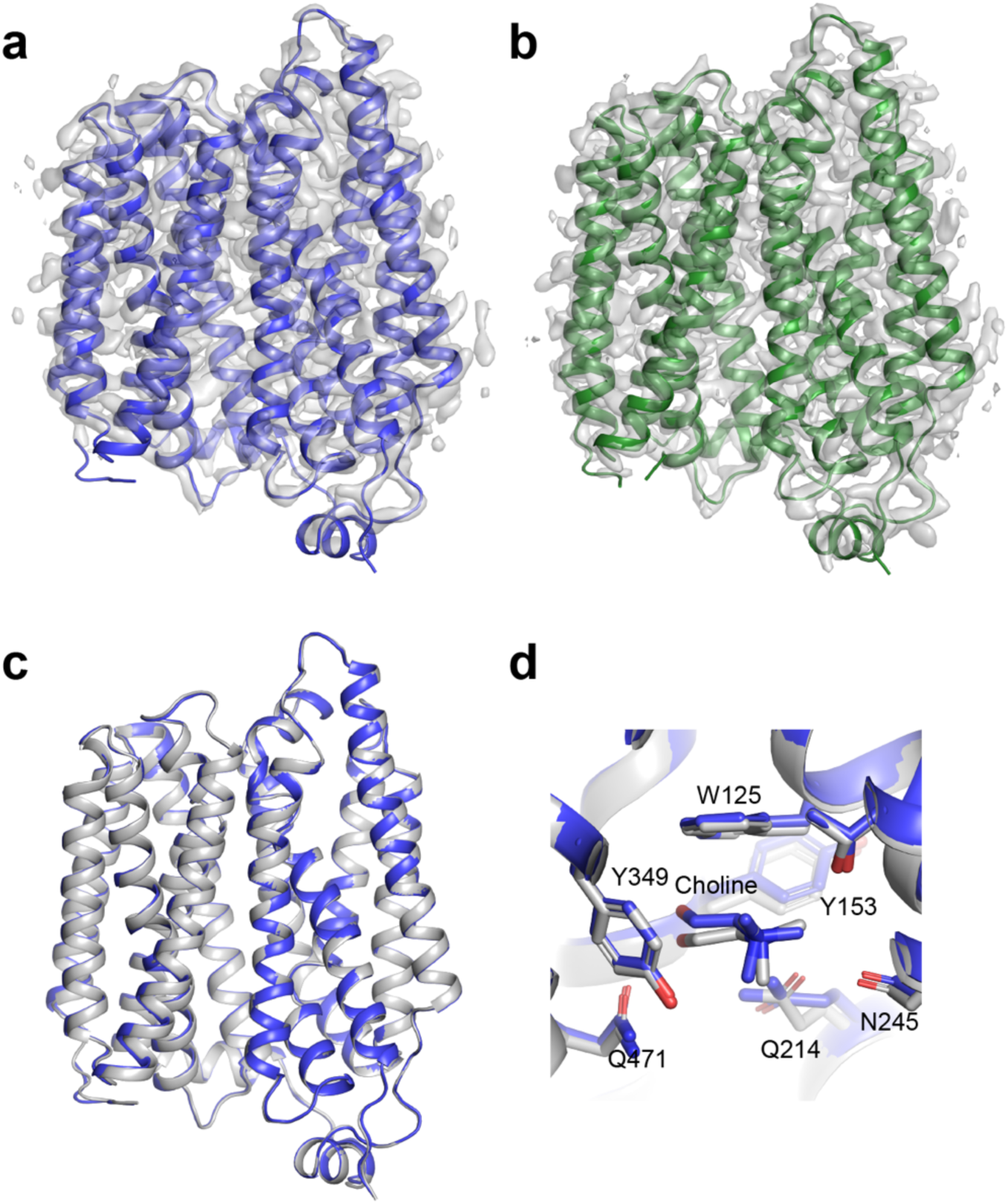
Cryo-EM structures of FLVCR1 determined in a substrate-free condition. **a-b**, Cryo-EM density maps and atomic models of FLVCR1 in endogenous choline (A) and endogenous ligand-bound (B) states. **c**, Superposition of choline-bound (grey) and endogenous choline-bound (blue) states. **d**, Superposition of substrate-binding sites in choline-bound (grey) and endogenous choline-bound (blue) states.

**Extended Data Fig. 6.**
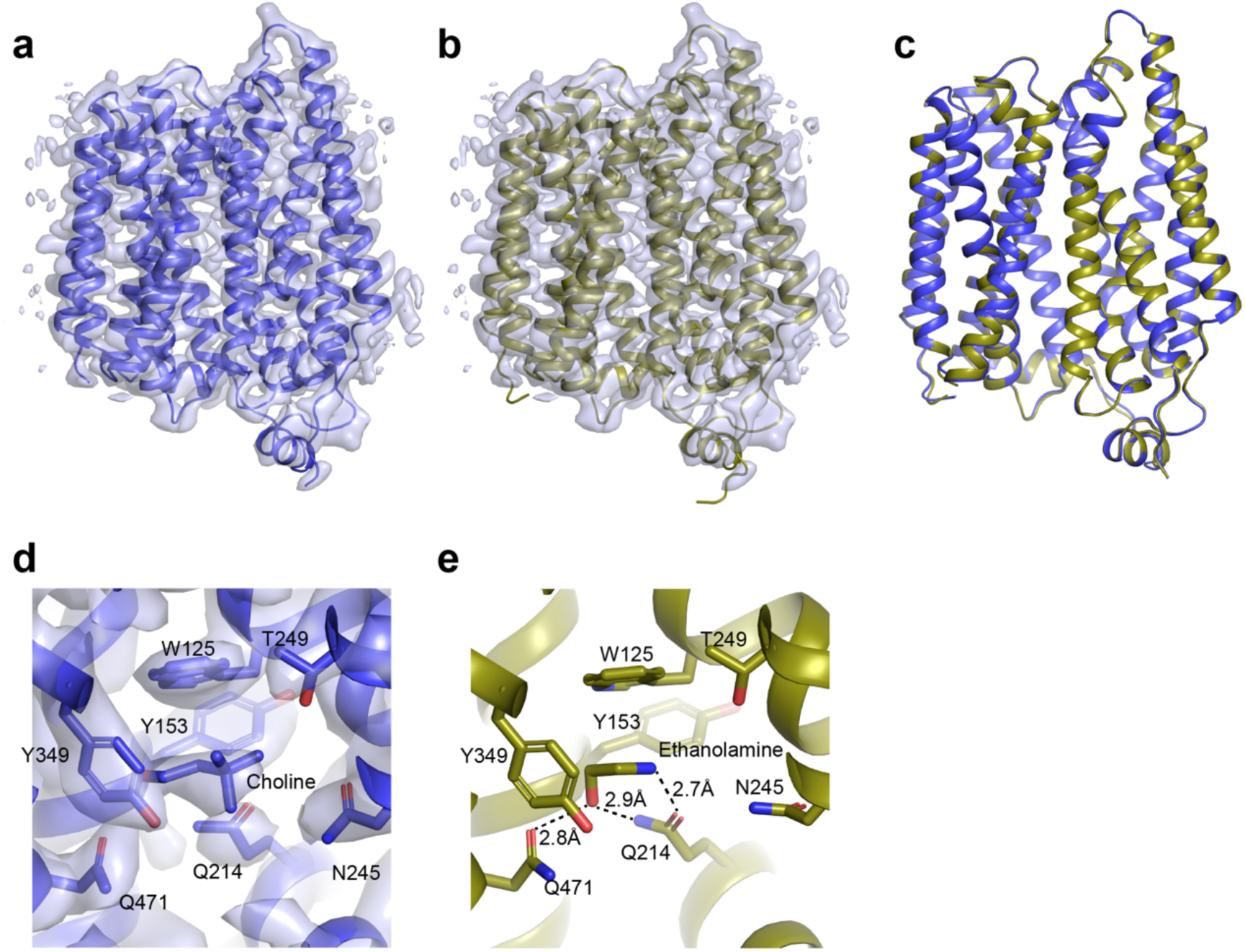
Cryo-EM structures of FLVCR1 determined in 1 mM ethanolamine. **a-b** Cryo-EM density maps of endogenous choline-bound FLVCR1 (a) and ethanolamine-bound FLVCR1 (b) states, determined from particles imaged in the presence of 1 mM ethanolamine. **c**, Superposition of ethanolamine-bound (gold) and choline-bound (blue) states. **d**, Substrate-binding site in endogenous choline-bound FLVCR1 state, determined from particles imaged in the presence of 1 mM ethanolamine. Residues and modelled substrates are shown as sticks. Density is shown as a grey isosurface and contoured at 3.0 σ. **e**, Coordination of ethanolamine in the substrate-binding site in ethanolamine-bound state.

**Extended Data Fig. 7.**
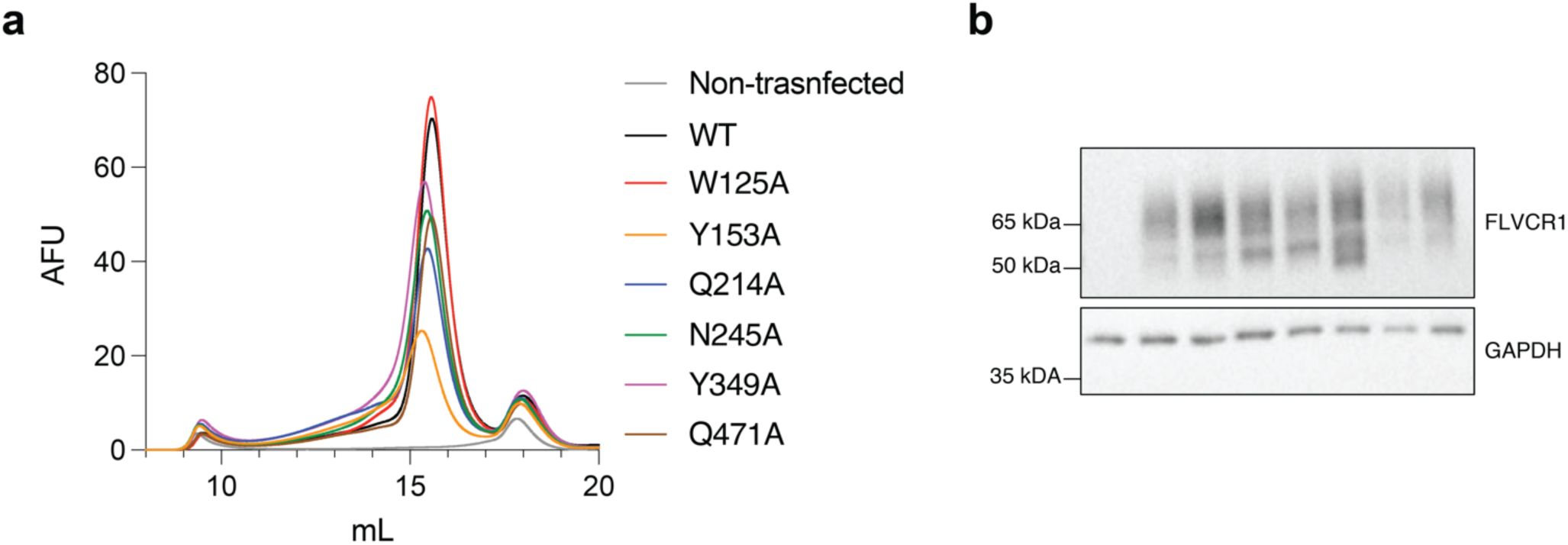
Expression and biochemical properties of FLVCR1 substrate-binding site mutants. **a**, FSEC analysis of wild-type or substrate-binding site mutants fused to mCerulean expressed in *FLVCR1*-knockout HEK293T cells. **b**, Western blot analysis of *FLVCR1*-knockout HEK293T cells expressing a vector control or wild-type or mutant *FLVCR1* cDNA. For western blot source data, see Supplementary Fig.1.

**Extended Data Fig. 8:**
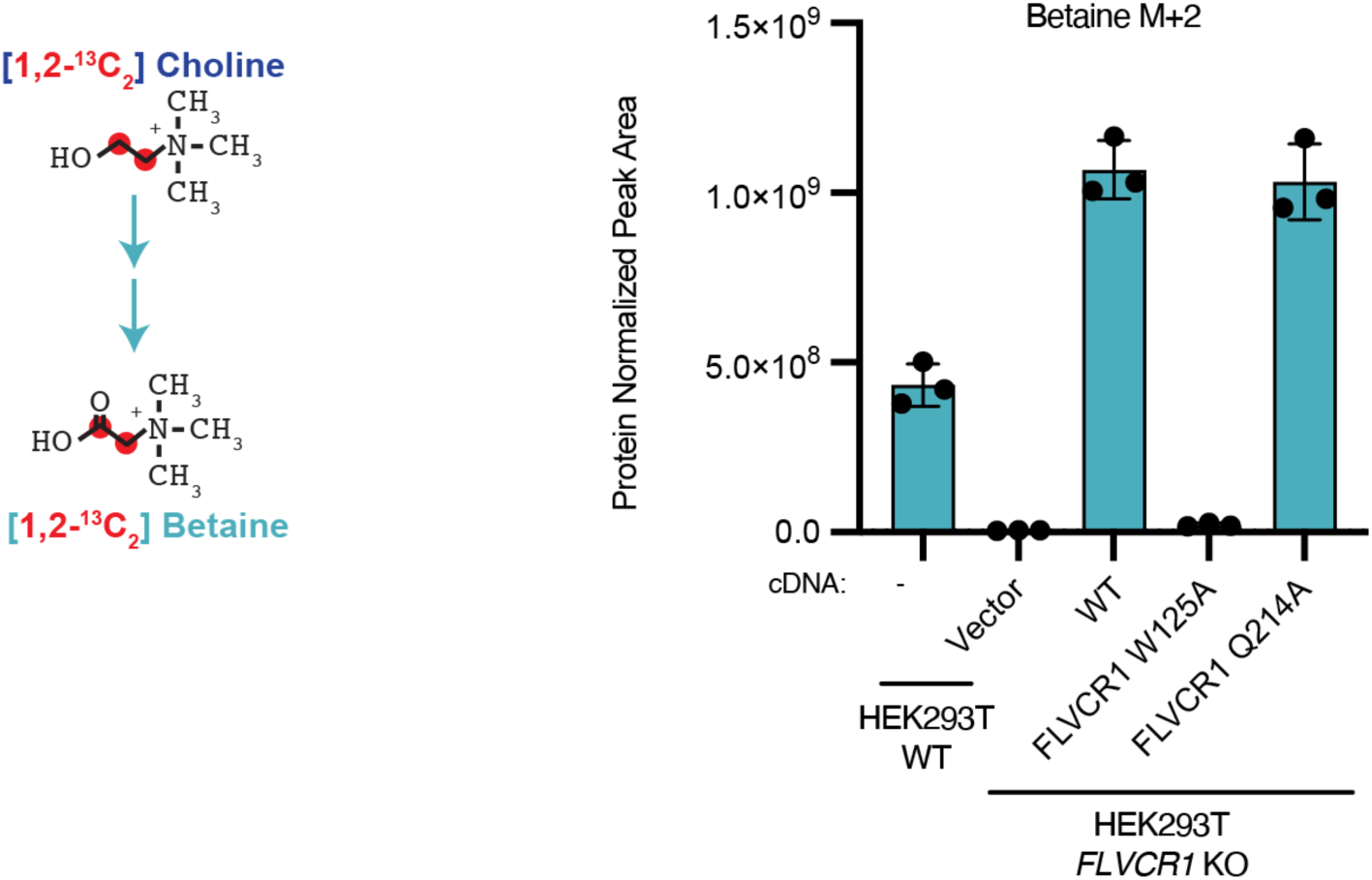
Incorporation of labeled choline into betaine requires FLVCR1. Schematic for tracing [1,2-^13^C_2_] choline into betaine (left) and the abundance of betaine M+2 after incubation with 21.5 µM [1,2-^13^C_2_] choline for 1 hour in *FLVCR1*-knockout HEK293T cells expressing a vector control or wild-type or mutant *FLVCR1* cDNA (right). Data shown as mean ± standard deviation; n=3.

## References

1. Vance, J. E. Phospholipid Synthesis and Transport in Mammalian Cells. Traffic 16, 1–18 (2015).

2. Patel, D. & Witt, S. N. Ethanolamine and Phosphatidylethanolamine: Partners in Health and Disease. Oxid. Med. Cell. Longev. 2017, (2017).

3. Kent, C. Phospholipid Metabolism in Mammals. Encycl. Biol. Chem. 3, 314–320 (2004).

4. Kennedy, E. P. Sailing to Byzantium. Annu. Rev. Biochem. 61, 1–28 (1992).

5. Gibellini, F. & Smith, T. K. The Kennedy pathway--De novo synthesis of phosphatidylethanolamine and phosphatidylcholine. IUBMB Life 62, 414–428 (2010).

6. Kennedy, E. P. SYNTHESIS OF PHOSPHATIDES IN ISOLATED MITOCHONDRIA: II. INCORPORATION OF CHOLINE INTO LECITHIN. J. Biol. Chem. 209, 525–535 (1954).

7. Yanatori, I., Yasui, Y., Miura, K. & Kishi, F. Mutations of FLVCR1 in posterior column ataxia and retinitis pigmentosa result in the loss of heme export activity. *Blood Cells*, Mol. Dis. 49, 60–66 (2012).

8. Rajadhyaksha, A. M. et al. Mutations in FLVCR1 cause posterior column ataxia and retinitis pigmentosa. Am. J. Hum. Genet. 87, 643–654 (2010).

9. Khan, A. A. & Quigley, J. G. Heme and FLVCR-related transporter families SLC48 and SLC49. Mol. Aspects Med. 34, 669–682 (2013).

10. Corbin, K. D. & Zeisel, S. H. Choline metabolism provides novel insights into nonalcoholic fatty liver disease and its progression. Curr. Opin. Gastroenterol. 28, 159–165 (2012).

11. Kenny, T. C. et al. Integrative genetic analysis identifies FLVCR1 as a plasma-membrane choline transporter in mammals. Cell Metab. (2023) doi:10.1016/j.cmet.2023.04.003.

12. Tsuchiya, M., Tachibana, N., Nagao, K., Tamura, T. & Hamachi, I. Organelle-selective click labeling coupled with flow cytometry allows pooled CRISPR screening of genes involved in phosphatidylcholine metabolism. Cell Metab. 35, 1072–1083.e9 (2023).

13. Kvarnung, M. et al. Mutations in FLVCR2 associated with Fowler syndrome and survival beyond infancy. Clin. Genet. 89, 99–103 (2016).

14. Meyer, E. et al. Mutations in FLVCR2 are associated with proliferative vasculopathy and hydranencephaly-hydrocephaly syndrome (Fowler syndrome). Am. J. Hum. Genet. 86, 471–478 (2010).

15. Lalonde, E. et al. Unexpected allelic heterogeneity and spectrum of mutations in Fowler syndrome revealed by next-generation exome sequencing. Hum. Mutat. 31, 918–923 (2010).

16. Drew, D., North, R. A., Nagarathinam, K. & Tanabe, M. Structures and General Transport Mechanisms by the Major Facilitator Superfamily (MFS). Chem. Rev. 121, 5289–5335 (2021).

17. Sauve, S., Williamson, J., Polasa, A. & Moradi, M. Ins and Outs of Rocker Switch Mechanism in Major Facilitator Superfamily of Transporters. Membranes (Basel). 13, (2023).

18. Yan, N. Structural Biology of the Major Facilitator Superfamily Transporters. Annu. Rev. Biophys. 44, 257–283 (2015).

19. Zhang, X. C., Zhao, Y., Heng, J. & Jiang, D. Energy coupling mechanisms of MFS transporters. Protein Sci. 24, 1560–1579 (2015).

20. Mödinger, Y., Schön, C., Wilhelm, M. & Hals, P.-A. Plasma Kinetics of Choline and Choline Metabolites After A Single Dose of SuperbaBoostTM Krill Oil or Choline Bitartrate in Healthy Volunteers. Nutrients vol. 11 (2019).

21. Garguilo, M. G. & Michael, A. C. Amperometric microsensors for monitoring choline in the extracellular fluid of brain. J. Neurosci. Methods 70, 73–82 (1996).

22. Brehm, R., Lindmar, R. & Löffelholz, K. Muscarinic mobilization of choline in rat brain in vivo as shown by the cerebral arterio-venous difference of choline. J. Neurochem. 48, 1480–1485 (1987).

23. Bianchi, L. et al. Extracellular levels of amino acids and choline in human high grade gliomas: an intraoperative microdialysis study. Neurochem. Res. 29, 325–334 (2004).

24. Plagemann, P. G. Choline metabolism and membrane formation in rat hepatoma cells grown in suspension culture. 3. Choline transport and uptake by simple diffusion and lack of direct exchange with phosphatidylcholine. J. Lipid Res. 12, 715–724 (1971).

25. Oswald, C. et al. Crystal structures of the choline/acetylcholine substrate-binding protein ChoX from Sinorhizobium meliloti in the liganded and unliganded-closed states. J. Biol. Chem. 283, 32848–32859 (2008).

26. Bärland, N. et al. Mechanistic basis of choline import involved in teichoic acids and lipopolysaccharide modification. Sci. Adv. 8, (2022).

27. Holm, L. Dali server: structural unification of protein families. Nucleic Acids Res. 50, W210– W215 (2022).

28. Xiao, Q. et al. Visualizing the nonlinear changes of a drug-proton antiporter from inward-open to occluded state. Biochem. Biophys. Res. Commun. 534, 272–278 (2021).

29. Leano, J. B. et al. Structures suggest a mechanism for energy coupling by a family of organic anion transporters. PLoS Biol. 17, e3000260 (2019).

30. Quigley, J. G. et al. Identification of a human heme exporter that is essential for erythropoiesis. Cell 118, 757–766 (2004).

31. Tsherniak, A. et al. Defining a Cancer Dependency Map. Cell 170, 564–576.e16 (2017).

32. Wainberg, M. et al. A genome-wide atlas of co-essential modules assigns function to uncharacterized genes. Nat. Genet. 53, 638–649 (2021).

33. Jackson, B. T. Identification of metabolic networks by genetic co-essentiality analysis. Nat. Rev. Mol. Cell Biol. 24, 378 (2023).

34. Taylor, A., Grapentine, S., Ichhpuniani, J. & Bakovic, M. Choline transporter-like proteins 1 and 2 are newly identified plasma membrane and mitochondrial ethanolamine transporters. J. Biol. Chem. 296, 100604 (2021).

35. Navale, A. M. & Paranjape, A. N. Glucose transporters: physiological and pathological roles. Biophys. Rev. 8, 5–9 (2016).

36. Goehring, A. et al. Screening and large-scale expression of membrane proteins in mammalian cells for structural studies. Nat. Protoc. 9, 2574–2585 (2014).

37. Mastronarde, D. N. Automated electron microscope tomography using robust prediction of specimen movements. J. Struct. Biol. 152, 36–51 (2005).

38. Suloway, C. et al. Fully automated, sequential tilt-series acquisition with Leginon. J. Struct. Biol. 167, 11–18 (2009).

39. Punjani, A., Rubinstein, J. L., Fleet, D. J. & Brubaker, M. A. cryoSPARC: algorithms for rapid unsupervised cryo-EM structure determination. Nat. Methods 14, 290–296 (2017).

40. Punjani, A., Zhang, H. & Fleet, D. J. Non-uniform refinement: adaptive regularization improves single-particle cryo-EM reconstruction. Nat. Methods 17, 1214–1221 (2020).

41. Scheres, S. H. W. Chapter Six – Processing of Structurally Heterogeneous Cryo-EM Data in RELION. in The Resolution Revolution: Recent Advances In cryoEM (ed. Crowther, R. A. B. T.-M. in E.) vol. 579 125–157 (Academic Press, 2016).

42. Terwilliger, T. C., Ludtke, S. J., Read, R. J., Adams, P. D. & Afonine, P. V. Improvement of cryo-EM maps by density modification. Nat. Methods 17, 923–927 (2020).

43. Jamali, K., Kimanius, D. & Scheres, S. H. W. A Graph Neural Network Approach to Automated Model Building in Cryo-EM Maps. in (2022).

44. Emsley, P., Lohkamp, B., Scott, W. G. & Cowtan, K. Features and development of {\it Coot}. Acta Crystallogr. Sect. D 66, 486–501 (2010).

45. Adams, P. D. et al. {\it PHENIX}: a comprehensive Python-based system for macromolecular structure solution. Acta Crystallogr. Sect. D 66, 213–221 (2010).

46. Pettersen, E. F. et al. UCSF Chimera—A visualization system for exploratory research and analysis. J. Comput. Chem. 25, 1605–1612 (2004).

47. Pettersen, E. F. et al. UCSF ChimeraX: Structure visualization for researchers, educators, and developers. Protein Sci. 30, 70–82 (2021).

48. Pavelka, A. et al. CAVER: Algorithms for Analyzing Dynamics of Tunnels in Macromolecules. IEEE/ACM Trans. Comput. Biol. Bioinforma. 13, 505–517 (2016).

49. Sievers, F. et al. Fast, scalable generation of high-quality protein multiple sequence alignments using Clustal Omega. Mol. Syst. Biol. 7, 539 (2011).

50. Jumper, J. et al. Highly accurate protein structure prediction with AlphaFold. Nature 596, 583– 589 (2021).

